# Quantitative bioactivity signatures of dietary supplements and natural products

**DOI:** 10.1101/2022.09.14.507630

**Authors:** Adam Yasgar, Danielle Bougie, Richard T. Eastman, Ruili Huang, Misha Itkin, Jennifer Kouznetsova, Caitlin Lynch, Crystal McKnight, Mitch Miller, Deborah K. Ngan, Tyler Peryea, Pranav Shah, Paul Shinn, Menghang Xia, Alexey V. Zakharov, Anton Simeonov

## Abstract

Dietary Supplements and Natural Products have minor oversight of their safety and efficacy. We assembled a collection of Dietary Supplements and Natural Products (DSNP) as well as Traditional Chinese Medicinal (TCM) Plant extracts, which were screened against an *in vitro* panel of assays, including a liver cytochrome p450 enzyme panel, CAR/PXR signaling pathways, and P-gp transporter assays, to assess their activity. This pipeline facilitated the interrogation of Natural Product-Drug Interaction (NaPDI) through prominent metabolizing pathways. In addition, we compared the activity profiles of the DSNP/TCM substances with those of an approved drug collection. Many of the approved drugs have well-annotated mechanisms of action (MOA) while the MOAs for most of the DSNP and TCM samples remain unknown. Based on the premise that compounds with similar activity profiles tend to share similar targets or MOA, we clustered the library activity profiles to identify overlap with the NCATS Pharmaceutical Collection to predict the MOAs of the DSNP/TCM substances. Overall, we highlight four significant bioactivity profiles (measured by p-values) as examples of this prediction. These results can be used as a starting point for further exploration on the toxicity potential and clinical relevance of these substances.

## Introduction

Small molecules and biologics are highly regulated for their pharmacological properties, safety, and efficacy. An approved drug, typically consisting of a single active pharmaceutical ingredient, can take 10 – 12 years and cost over a billion dollars to develop.^1^ In stark contrast, The Dietary Supplement Health and Education Act of 1994 (DSHEA) regulates dietary supplements for Good Manufacturing Practice (GMP), but it does not standardize quality, safety, or efficacy of these compounds.^2^ While facilitating easier consumer access, it places the onus of quality, safety, and efficacy on consumers and regulators rather than the company that manufactured the product. This leads to serious concerns when health claims are made by manufacturers with no supporting evidence and is exacerbated by the lack of available safety data, especially for substances derived from plants or botanicals that contain a number of ingredients often of unknown sourcing.^3^

Since the DSHEA enactment, there has been an unfortunate proliferation of unsubstantiated claims of healing combined with multiple instances of substances “pulled from the shelves” due to contamination and/or adulteration, leading to serious health incidents. This began almost immediately in the 1990s with Ephedra^4^, a plant extract preparation with methamphetamine-like components that was used in weight-loss supplements that was ultimately banned for sales by the FDA in 2004, and continues to the present day with Kratom^5^, a plant being sold to manage pain and opioid-withdrawal that was removed from the market because of apparent properties that expose users to the risks of addiction, abuse, and dependence. A recent case involved belladonna, a plant used in baby teething tablets due to its analgesic properties, was recalled in 2010 and reformulated in 2011. However, even with these steps taken, it was later linked to the deaths of 10 children in 2017 and subsequently recalled.^6^ These adverse events were recently summarized by the FDA, with more than 25,000 incidents reported for dietary supplements between the years 2004-2016^7^ (contributing up to 20,000 ER visits yearly^8^), leading to the removal of hundreds of products from store shelves.^9^

One avenue of research into testing the toxicity and efficacy of these substances involves the study of drug-drug interactions, or in this case, natural product-drug interactions (NaPDI).^10^ Natural products include materials derived from animals, herbals, macroscopic fungi, and bacteria as well as materials derived from them (e.g. vitamins, amino acids, etc.).^11^ A classic case is grapefruit juice, a CYP3A4 inhibitor that metabolizes more than 85 drugs.^12^ Another natural product example is Vitamin K, which is involved in blood clotting and decreases the effectiveness of the blood thinner warfarin and similar drugs.^13^

In an effort to enhance proactive research efforts in the context of natural product toxicity evaluation and NaPDI^14^, we assembled a collection of commercially available DSNP along with the National Cancer Institute Traditional Chinese Medicine (NCI TCM) collection to characterize these substances against a panel of biochemical and cellular assays to assess their impact on drug metabolism and transport pathways. These results were then compared to those obtained from similar profiling of an approved drugs collection through clustering and PCA approaches. The results derived from this study serve as a starting point for further exploration on the toxicity potential and clinical relevance of these substances.

## Results

For this endeavor, we used two discrete compound collections. The first, the NCI Library of Traditional Chinese Medicine (TCM) Plant Extracts, kindly provided by the Natural Products Branch of the Developmental Therapeutics Program in the Division of Cancer Treatment and Diagnosis at the NCI. The second library, assembled by us for the purpose of the study, was the Dietary Supplement and Natural Product (DSNP) library, a carefully curated collection of commercially sourced dietary supplements and natural products. Substances were primarily chosen based on sales revenue^15^ and reported use^16^ to capture those impacting the largest population (see Methods section). All substances were tested in quantitative high-throughput screening (qHTS) format. Screening in qHTS format yielded concentration-response-curves (CRC) from the primary screen, enabling us to prioritize active compounds (**Figure S1**). We employed the following assays against the assembled libraries to assess for bioactivity: liver cytochrome p450 enzyme panel, CAR/PXR signaling pathways, and P-gp transporter assays, which represent prominent pathways associated with drug metabolism, side-effects, and disposition.

### P450 assays

P450 activity is an essential part of any preclinical drug package to inform about potential drug-drug interactions (DDI), or in our case natural product-drug interactions (NaPDI), which have been previously described.^17-19^ To enable qHTS, we employed a P450-Glo assay (Promega, Madison, WI) based on previous work performed at NCATS and HTS-amenability. We screened a total of 964 samples in qHTS format, made up of the DSNP (300 samples) and TCM (664 samples) libraries, against five isoforms of cytochrome P450 (CYP1A2, CYP2C9, CYP2C19, CYP2D6, and CYP3A4). The assay performance was excellent with Z’-factors ranging from 0.68 to 0.92 (**Tables S2a**).^20^ The intraplate controls Furafylline (CYP1A2), Sulfaphenazole (CYP2C9), Ketoconazole (CYP2C19), Quinidine (CYP2D6), and Ketoconazole (CYP3A4) also performed well, with mean IC_50_ values [µM] of 2.60 (Minimum Significant Ratio; MSR = 3.57), 0.04 (MSR = 1.78), 5.17 (MSR = 1.82), 0.10 (MSR = 5.98), and 0.01 (MSR = 1.18), respectively (**Figure S2 and Table S1c**), which is in agreement with previously published values.^21-24^

Assay responses were evaluated as previously described.^21,25-27^ Selectivity was defined as being active towards the isozyme of interest and inactive against the other CYPs and/or exhibiting a >5-fold IC_50_ ratio if activity was observed. **Table 1** displays the breakdown of the number of actives and potency ranges for each CYP tested against the DSNP and TCM libraries. Overall, a broad-spectrum of inhibitory activity for all five isozymes were observed for both the DSNP and TCM collections, with 8 – 56% and 10 – 91%, respectively, inhibiting any one CYP isozyme (**Table 1**).

**Table 1.**
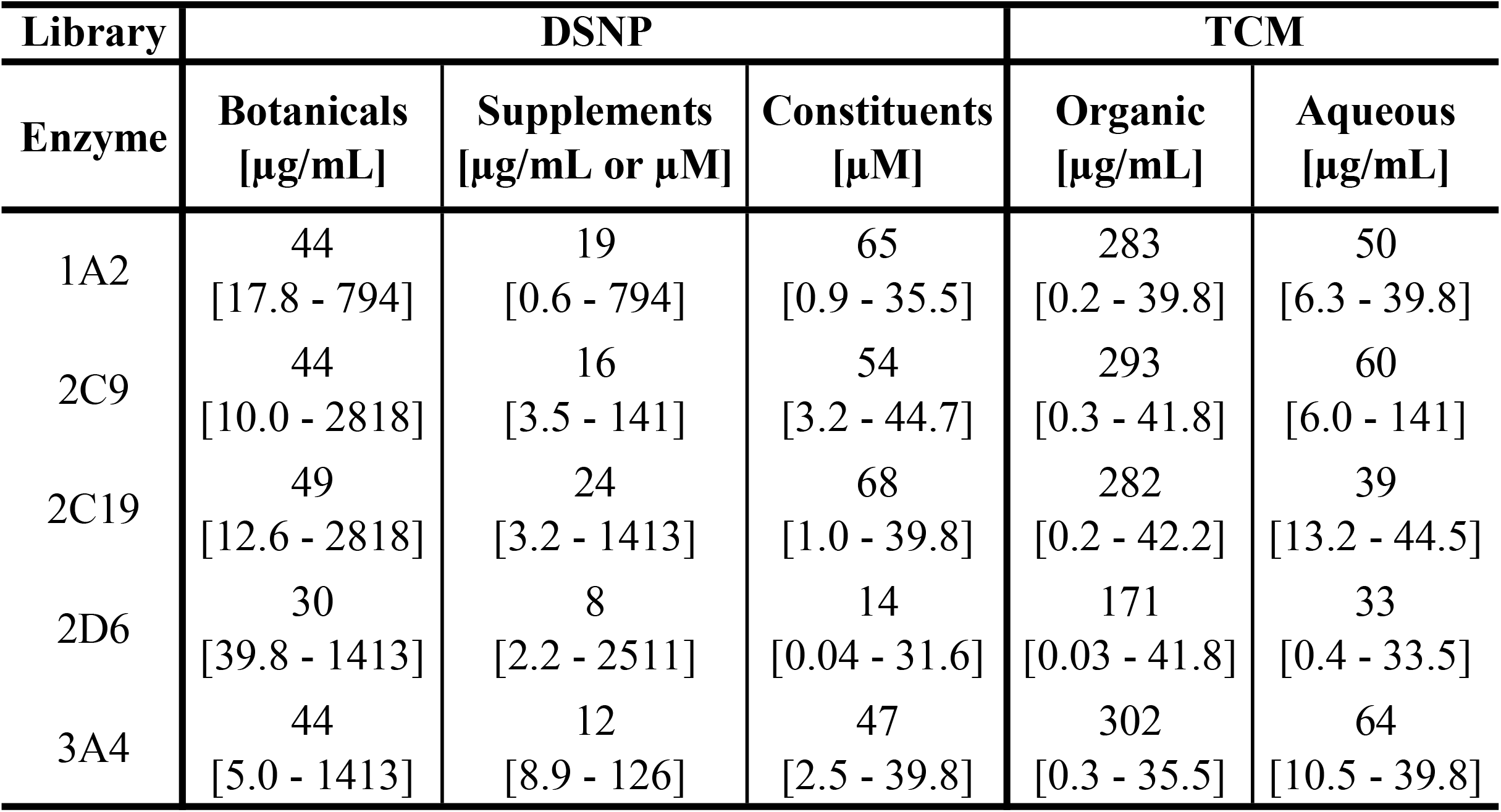
P450 active breakdown.

| **Library** | **DSNP** |  |  | **TCM** |  |
| --- | --- | --- | --- | --- | --- |
| **Enzyme** | **Botanicals**<br>**[μg/mL]** | **Supplements**<br>**[μg/mL or μM]** | **Constituents**<br>**[μM]** | **Organic**<br>**[μg/mL]** | **Aqueous**<br>**[μg/mL]** |
| 1A2 | 44<br>[17.8 - 794] | 19<br>[0.6 - 794] | 65<br>[0.9 - 35.5] | 283<br>[0.2 - 39.8] | 50<br>[6.3 - 39.8] |
| 2C9 | 44<br>[10.0 - 2818] | 16<br>[3.5 - 141] | 54<br>[3.2 - 44.7] | 293<br>[0.3 - 41.8] | 60<br>[6.0 - 141] |
| 2C19 | 49<br>[12.6 - 2818] | 24<br>[3.2 - 1413] | 68<br>[1.0 - 39.8] | 282<br>[0.2 - 42.2] | 39<br>[13.2 - 44.5] |
| 2D6 | 30<br>[39.8 - 1413] | 8<br>[2.2 - 2511] | 14<br>[0.04 - 31.6] | 171<br>[0.03 - 41.8] | 33<br>[0.4 - 33.5] |
| 3A4 | 44<br>[5.0 - 1413] | 12<br>[8.9 - 126] | 47<br>[2.5 - 39.8] | 302<br>[0.3 - 35.5] | 64<br>[10.5 - 39.8] |

For the DSNP collection, a wide range of P450 activity was observed, with 12% of substances pan-active against all five isozymes while 36% of substances were inactive (**Figure 1)**. The remaining substances were distributed evenly among the combination of activity in one to four CYPs, with a range of 8 –18%. A total of 88 botanicals, 43 supplements, and 169 constituents were separated into subgroups of the DSNP library and CYP inhibition was calculated for each category. (**Tables 1 and S2a**). For all three categories, CYP2D6 consistently exhibited the lowest number of actives, followed by CYP3A4, CYP2C9, CYP1A2, and CYP2C19, respectively. Overall, 64% of these substances have the potential to impact at least one of the major human P450 metabolizing enzymes.

**Figure 1.**
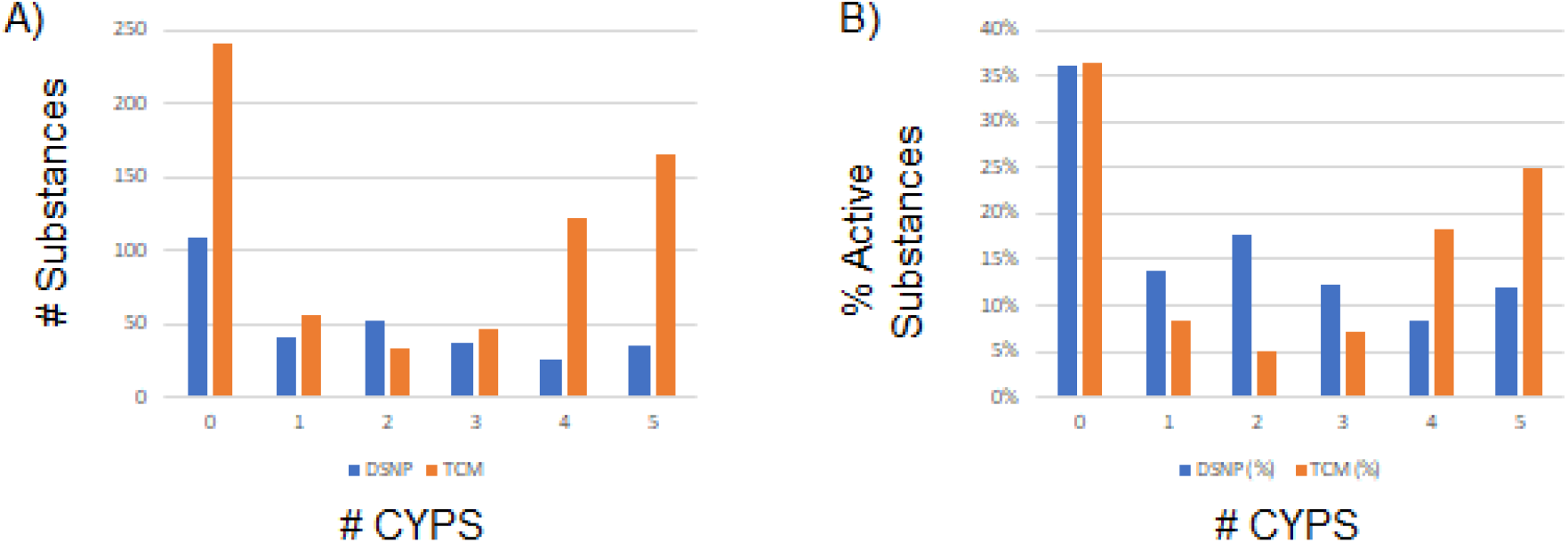
Distribution and differences in CYP activity between DSNP versus TCM libraries. (a) Distribution of DSNP (blue) and TCM (orange) compounds that were active against a given number of CYP isozymes.

Pan-activity, an undesirable characteristic due to potential NaDPI, was observed in 36 samples (12%), with the majority (23 samples) belonging to the botanicals subset. While there are no potency cutoffs for what would equate to an NaDPI versus a specific CYP, substances that merit further study include Angelica Root Oil (NMIX00599544) and Rhubarb Fluid Extract (NMIX00599581), two botanicals that exhibited the strongest pan-activities (**Table 2**). Two other substances met these criteria, the constituents Isorhamnetin (NCGC00163572; IC_50_ range of 4.5 – 11.2 µM) and Pentagalloyl glucose (NCGC00180839; IC_50_ range of 1.0 – 14.1 µM). Their respective parent substances, Boldo Leaf (NMIX00599541) and Peony Root (NMIX00599575), exhibited pan-activity (IC_50_ range of 39.8 – 126 µg/mL) and inactivity, respectively, with the latter result being somewhat surprising given the strong activity observed from Pentagalloyl glucose. By including constituents in the library, parent and substance activity can be easily compared. Separately, inactivity against all the isoforms was observed within each group of the DSNP library, with 25 botanicals (28%), 13 supplements (30%), and 70 constituents (41%) exhibiting no activity, totaling to 108 substances (36%).

**Table 2.**
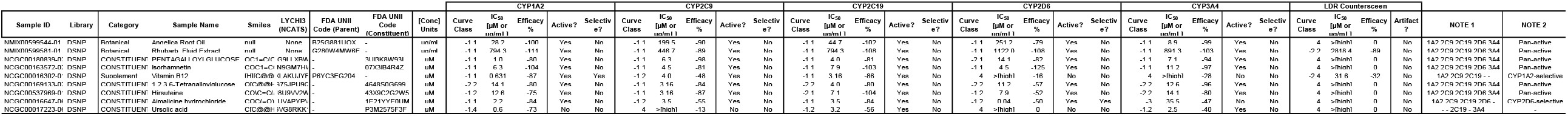

The most potent substances against each isoform were predominantly constituents. Ursolic acid (NCGC00017223) is a component of Holy Basil (NMIX00601615). In this case, both the constituent and the parent substance were potent against CYP3A4, with Urosolic acid having an IC_50_ value of 2.5 μM and Holy Basil with an IC_50_ of 44.7 μg/mL, though the parent substance was shown to have pan-activity. 1,2,3,6-Tetragalloylglucose (NCGC00169133), a constituent of Peony Root (NMIX00599575), exhibited activity against CYP2C9 with an IC_50_ of 3.2 μM. In contrast, Peony Root had no observable activity. Hirsuteine (NCGC00537969) is found in Cat’s Claw (NMIX00601608; previously identified as a CYP3A4 inhibitor^28^) and was highly potent against CYP2C9 with an IC_50_ of 3.2 μM. In contrast, Cat’s Claw was highly active, with activity seen against all isoforms. Ginesenoside RB1 (NCGC00385315) was most potent against CYP2C19 at an IC_50_ of 1 μM and is found in Ginseng (NMIX00599534 and NMIX00601612). Both Ginseng samples were active only against CYP2C9. As observed in a previous study, Ajmalicine (NCGC00016647) was potent against CYP2D6 with an IC_50_ = 0.04 μM,^29^ and it has been recognized as a component from the above mentioned Cat’s Claw. Lastly, the only non-constituent in this list, Vitamin B12 (NCGC00016302), was the most potent against CYP1A2 at an IC_50_ of 0.631 μM.

The largest number of constituents associated with a mixture occurred with Saw Palmetto (NMIX00599537) and Peony Root Powder, each composed of 18 substances (**Table S2b**). For Saw Palmetto **(Figure S3**), which itself was pan-active with an IC_50_ range of 22.4 – 79.4 µg/mL, three compounds, Linoleic acid (NCGC00344326), Oleic acid (NCGC00344330), and Lauric acid (NCGC00090919), appear to be responsible for the majority of its P450 activity except for CYP2D6, where none of the 18 samples were active. While nine of the samples displayed some inhibitory activity, the remaining nine samples were inactive against all CYPs tested. In contrast, the Peony Root mixture (NMIX00599575) was inactive against all CYPs over the concentration range tested, contrary to the activity observed in its components, where 2 of the 18 constituents, Pentagalloyl glucose (NCGC00180839) and NP-003686 (NCGC00169133), were pan-active. Similar to Saw Palmetto, 7 of the 18 constituents were completely inactive, while the remaining 11 constituents had varying degrees of CYP inhibition.

A similar analysis was performed on the NCI TCM collection, which consisted of 664 samples (132 species and 332 extracts) that were treated with either aqueous or organic solvents. We first chose to compare the two extraction types, as this is an important part of the process when taking TCM. **Table 1** compares the overall performance between the two extraction types; of the 332 organic extracts, 52 – 91% inhibited any one CYP isozyme, with four of the CYPs at >85%, along with similar ranges in potency (**Tables 1 and S2a**). Of the 332 organic extract samples, 153 (46%) exhibited pan-activity, covering 79 (60%) species. Only 17 (5%) of organic extract samples did not exhibit any activity. From the actives group, 61 to 131 (18% – 39%) samples were of our highest quality CRC’s (−1.1), for CYP1A2, CYP2C9, CYP2C19, and CYP3A4, with strong inhibition observed by all except for CYP2D6. This is in sharp contrast with the aqueous extracts, with only 10 – 19% of samples inhibiting any one CYP isozyme, along with weaker potencies, with aqueous extract potencies 13- to 66-fold weaker than their organic counterparts. Of the 332 aqueous extracts, 224 (67%) extracts were inactive or 76 out of132 species (58%). Overall, ∼5-fold less activity was observed in the aqueous extract when compared to organic. Importantly, this indicates that the activity is largely being derived from the organic extracts.

Continuing our analysis of the TCM collection, a wide range of P450 activity was observed with 25% of the samples pan-active against all five isozymes, in contrast to 36% that were inactive (**Figure 1)**. The remaining substances demonstrated a modest increase in activity (5 – 18%) compared to the number of substances that were active against one to four CYPs. Upon closer examination of the relationship between extract activity and the species they were derived from, 423 (64%) samples from 128 (97%) different species were found to inhibit any one CYP isozyme (**Tables 1, S2a, and S2c**). Due to the high activity observed, we applied a filter to identify species with ≥6 samples which resulted in 88 species, and this larger set of samples from a species might provide more insight to its bioactivity against P450. As shown in **Table S2c**, 79 out of 88 samples (90%) had ≥50% of their extract show activity against any CYP isoform, with 19 species having all samples exhibit inhibitory activity against any one CYP isozyme., A total of 165 samples from 24 (27%) species had ≥50% of their extracts demonstrate pan-CYP activity, exemplifying the broad *in vitro* P450 activity of this collection **(Tables S2c and S2d)**. A majority of this activity arose from 153 (93%) organic extracts. Only one species, *Lindera aggregata* saw pan-activity among all six of its organic and aqueous samples. One additional species, *Rosa rugosa*, had more than one aqueous extract active among its group of extracts. Overall, 64% of the mixtures tested have a role in one of the major human P450 metabolizing enzymes.

The most potent samples consisted entirely of organic extracts (IC_50_ values < 0.3 μg/mL). CYPs 1A2, 2C9, 2C19, 2D6, and 3A4 were derived from *Angelica dahurica* (root of the Holy Ghost, NMIX00590230), *Sophora flavescens* (Ku Shen, NMIX00590416), *Juncus effusus* (Soft Rush, NMIX00590400), *Corydalis yanhusuo* (NMIX00590373) and *Piper nigrum* (Black pepper, NMIX00590313), respectively (**Figure S4** and **Table S2a**). The four species that exhibited no activity among all samples tested were *Codonopsis pilosula* (poor man’s ginseng), *Dimocarpus longan* (dragon’s eye), *Ganoderma lucidum* (reishi mushroom), and *Lycium barbarum* (Goji berry).

Next, we compared the CYP activity between the two collections. As shown in **Figure 1**, an equal proportion (36%) of the DSNP and TCM libraries were inactive against all five isozymes. Pan-activity was increased two-fold in the TCM library when compared to the DSNP library (25% and 12%, respectively). While the DSNP library percent activity was flat when comparing activity for single or multiple isoforms, there was a 2- to 3-fold increase observed in the TCM library for extracts with >3 isozymes. We identified the previously reported and well-characterized CYP2D6-selective substance Berberine (IC_50_ = 2.2 μM) that is the main component of Goldenseal, a popular supplement promoted for the treatment of multiple ailments.^30^ The supplements Cannabidiol (CBD; NCGC00386518) and Omega-3 fatty acids (NMIX00599569) were pan-active, with IC_50_ values ranging from 7.9 – 14.1 µM and 100 – 158 µg/mL, respectively, and these observations were generally in agreement with previous findings.^31,32^ In contrast, Capsicum (NMIX00599568) a potent botanical that exhibited pan-activity with IC_50_ values < 50 µg/mL, was found to have little to no activity in microsomes from hepatocyte primary cultures.^33^

Finally, results were analyzed for P450 activation, which was infrequently observed. CYP isozyme activation, or the increased rate of proluciferin conversion, is typically substrate dependent, so the present assays would not comprehensively characterize this interaction. Nonetheless, 37 (∼4%) samples from both collections exhibited P450 activation for CYP1A2 (1 sample), CYP2C9 (8 samples), CYP2C19 (28 samples), CYP2D6 (3 sample), and CYP3A4 (3 samples, **Table S2f**). 28 out of 37 samples exhibited activation in CYP2C19, while only 5 out of 37 came from the TCM Library. Of these actives, a majority showed weak activity, with only six samples exhibiting strong CRCs (1.1. or 2.1). Samples that induced CYP2C9 activation were Formononetin (NCGC00017269; 3.2 μM), Gingerol (NCGC00163131; 11.2 μM), and an aqueous extract from *Pueraria lobata* (kudzu, NMIX00590752; 3.8 μg/mL). Similarly, aqueous extracts of *Panax notoginseng* (Chinese ginseng, NMIX00590550; 0.5 μg/mL) and *Verbena officinalis* (NMIX00590580; 0.53 μg/mL) activated CYP2C19, while the constituent Isosteviol (NCGC00485884; 25.1 μM) activated CYP3A4.

### CAR and PXR reporter gene assays

Modulation of CAR and PXR have been shown to cause up/down regulation of CYP enzymes at the mRNA, protein, and activity levels. To this end, we profiled both collections for potential CAR and/or PXR activity, employing a previously published 1536-well qHTS assay.^26,34^ The overall assay performance was excellent for both receptors and modes of action tested. Mean Z’-factors for the agonist assays of 0.51 and 0.70 were calculated for CAR and PXR, respectively (**Table S1b**). The intraplate positive controls for the agonist assays, CITCO and Rifampicin, performed as expected, with mean EC_50_ values of 0.84 µM (MSR = 3.0) and 19.4 µM (MSR = 1.6) for CAR and PXR, respectively. Mean Z’-factors of 0.68 and 0.58 for CAR and PXR, respectively, were noted for the antagonist assays. The intraplate positive controls for the antagonist assays, PK11195 and SPA70, performed as expected, with mean IC_50_ values of 4.9 µM (MSR = 1.6) and 0.73 µM (MSR = 3.6), for CAR and PXR, respectively. For the antagonist cytotoxicity assays, the mean Z’-factors of CAR and PXR were 0.83 and 0.79, respectively. The intraplate positive control Tetraoctyl ammonium bromide performed as expected, with mean IC_50_ values of 68.2 µM (MSR = 1.2) and 49.7 µM (MSR = 1.4), for CAR and PXR, respectively.

For the CAR assays, 141 agonists (15%) and 69 antagonists (7%) were identified as active from both collections (**Figure 2a and Table S3a; see Methods section**). For the DSNP library, 37 agonists (12% of the collection; 10 botanical, 8 supplements, 19 constituents) and 68 antagonists (23% of the collection; 18 botanical, 4 supplements, 46 constituents) exhibited activity. To highlight the strongest agonists, we first applied potency and efficacy triage filters of < 15 μM or 75 µg/mL and > 40%, respectively, then removed weaker CRCs of 1.2 and 2.2. For antagonists, we applied the same potency filter with an increased efficacy cutoff of ≤ -50% efficacy and inactive in the cytotoxicity assay. Application of the filters resulted in 10 agonists (one supplement and nine constituents) and two antagonists (both constituents) (**Table 3**). The one supplement agonist, Tryptophan (NCGC00015994), exhibited an EC_50_ value of 3.98 µM while the nine constituent agonists displayed an EC_50_ range of 3.98 µM – 14.1 µM. Also of interest were four constituent compounds, starting with the two isoflavonoid glucosides Daidzin (NCGC00163532) and Ononin (NCGC00169054); both exhibited strong CAR agonist activity and were found to be linked to the parent substance Astragalus (NMIX00599594) which was inactive. This contrasts with a previous publication indicating Astragalus induced PXR activity.^35^ The other two related compounds with strong responses were Demethoxyyangonin (NCGC00091904) and Yangonin (NCGC00091909). Both are linked to Kava Kava Root (NMIX00599602), which had an EC_50_ value of 141 μg/mL, corroborating previous observations and serving as a possible example of individual components exhibiting similar activity to that of the parent mixture.^36^ The two CAR antagonists identified were the structurally similar constituents Isocorynoxeine (NCGC00482763) and Corynoxine B (NCGC00482767), with IC_50_ values of 3.98 µM each. Both are found in Cat’s Claw (NMIX00601608), which was inactive in all nuclear receptor modes.

**Table 3.**
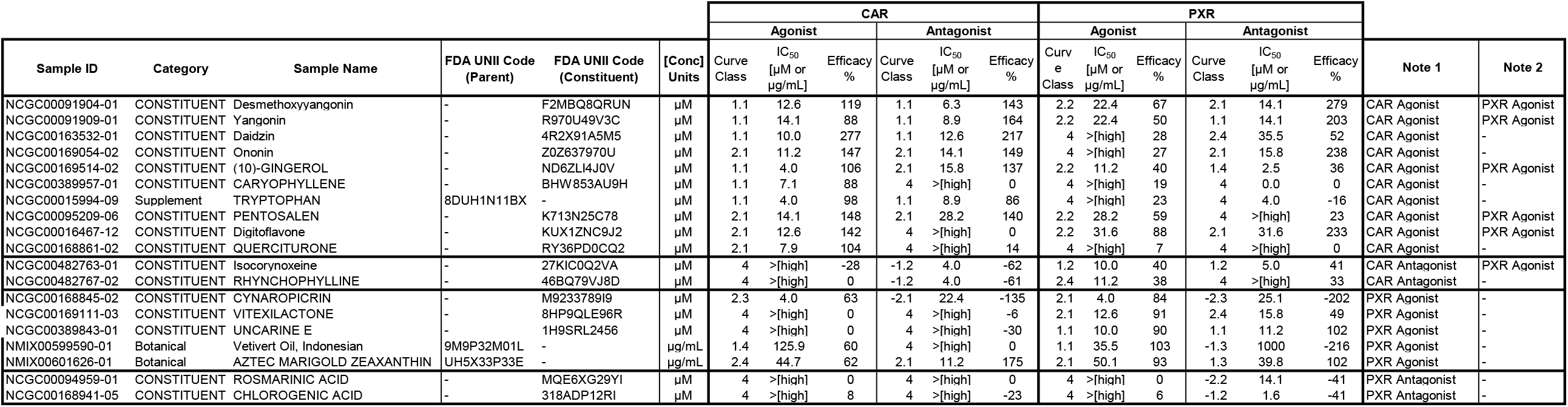

**Figure 2.**
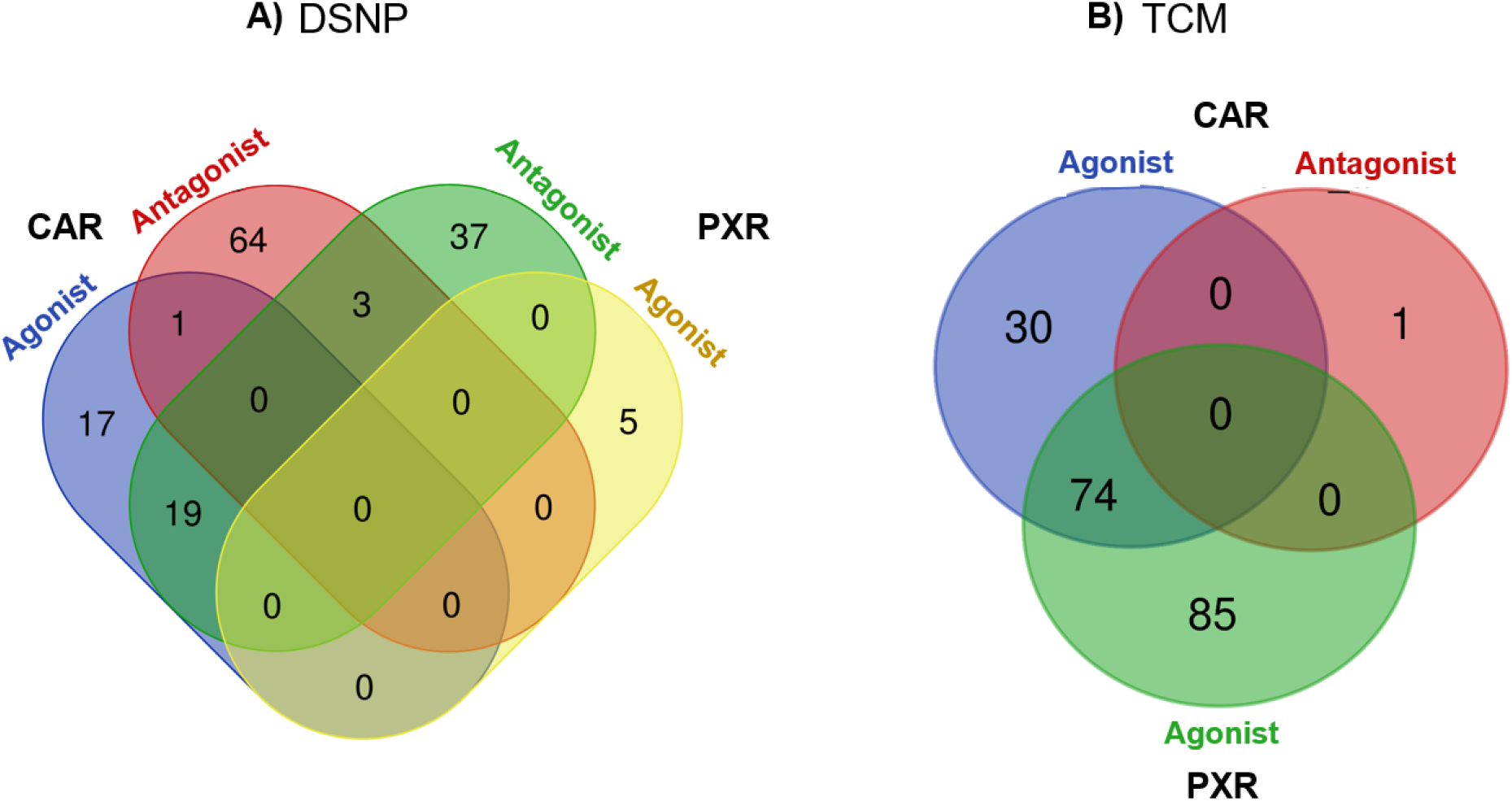
CAR and PXR agonist and antagonist actives in the DSNP (Panel A) and TCM (Panel B) collections.

In the NCI TCM library, we observed 104 CAR agonists (16%) that covered 50 different species (made up of 90 organic and 14 aqueous extracts; **Figure 2b**). The only CAR antagonist identified was derived from an aqueous extract of *Polygala tenuifolia* (yuan zhi; NMIX00590780; **Table 4**), exhibited an IC_50_ value of 22.7 µg/mL, but was weak with a CRC of 2.2 and an efficacy of -54%. For the CAR agonists, we applied the same filters mentioned above, leaving 45 agonists (23 species, 42 organic, 3 aqueous; **Tables S4a**). Due to the high number of agonist actives, we applied an additional filter, identifying species with ≥ 3 active samples, leaving five species of three samples each (15 samples total; all organic extracts): *Angelica dahurica* (root of the Holy Ghost), *Alpinia officinarum* (lesser galangal), *Dictamnus dasycarpus, Evodia rutaecarpa* (Fruits of Evodia, Wu Zhu Yu), and *Alpinia oxyphylla*, with an EC_50_ range of 11.3 to 35.7 µg/mL (**Table 4**). The most potent agonist in the NCI TCM collection was an organic extract of *Cynanchum atratum* (Bai wei, NMIX00590250) at 6.4 μg/mL.

**Table 4.**
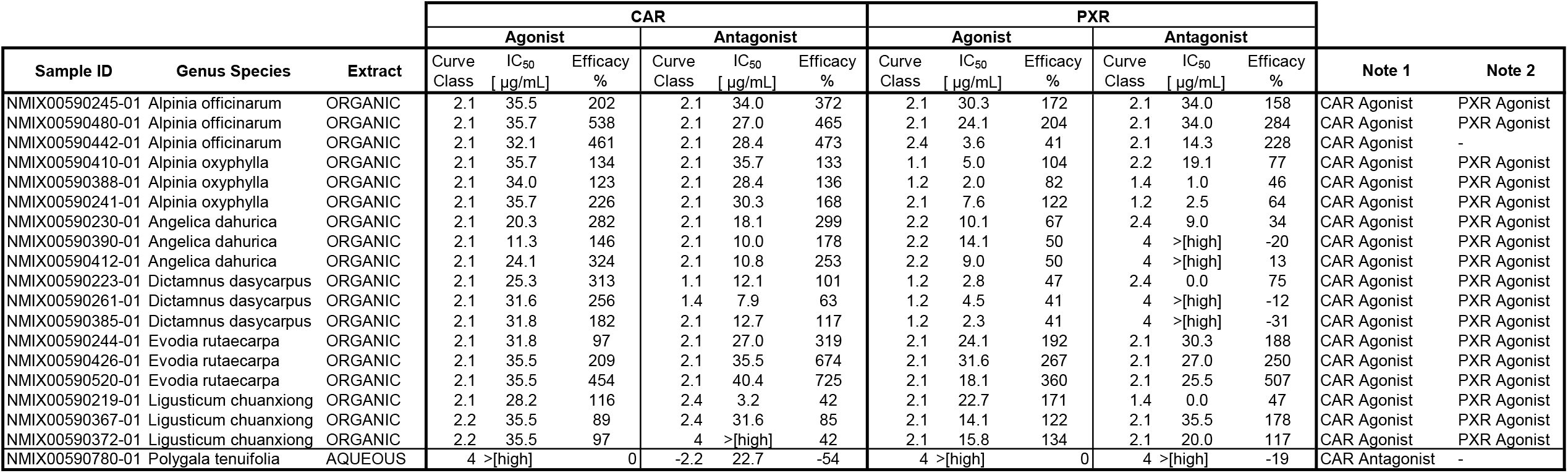
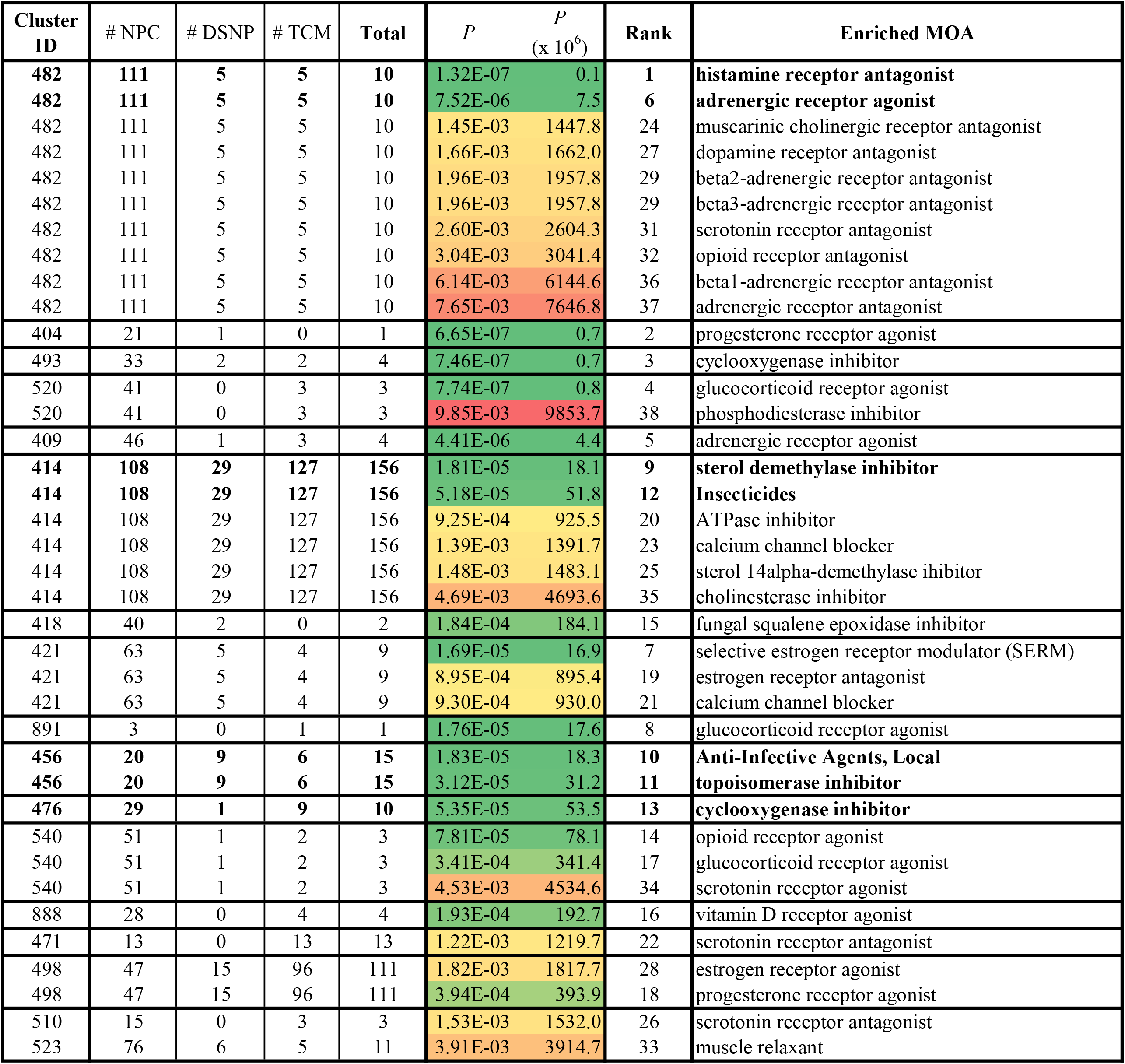

Overall, 219 agonists (23%) and 5 antagonists (0.5%) were initially identified for the PXR assays (**Table S3b)**. The DSNP library contained 59 agonists (20%; 20 botanical, 11 supplements, 28 constituents) as well as five antagonists (2%; one botanical, two supplements, two constituents). Applying criteria filters to the agonist actives, five agonists remained (two botanicals and three constituents; **Table 3**). The two botanical substances Vetivert Oil (Indonesian; NMIX00599590) and Aztec Marigold Zeaxnthin (NMIX00601626) displayed EC_50_ values of 35.5 μg/mL and 50.1 µg/mL, respectively. The three constituents, Cynaropicrin (NCGC00168845; the most potent), Uuncarine E (NCGC00389843), and Vitexilactone (NCGC00169111), had EC_50_ values of 10.0 µM, 12.6 μM, and 3.98 μM, respectively. The antagonists Chlorogenic acid (NCGC00168941) and Rosmarinic acid (NCGC00094959) exhibited EC_50_ values at 1.6 μM and 14.1 μM, respectively.

The NCI TCM library yielded 160 PXR agonists (24%; 77 species; 150 organic, 10 aqueous; 45% of organic extracts; 63% of species tested) and no antagonists. Applying the above agonist filters, 36 compounds remained (24 species, 36 organic, 0 aqueous). Due to the high number of agonist actives, we applied an additional filter, identifying species with ≥ 3 active samples, leaving two species of three samples each (6 samples total; all organic extracts): *Ligusticum chuanxiong* and *Evodia rutaecarpa* (Fruits of Evodia, Wu Zhu Yu), with EC_50_ ranges of 14.1 – 31.6 µg/mL, highlighted in **Table 4**.

Overlapping receptor and/or mode of activity was observed in both libraries using our final active criteria. For the DSNP library, 6 out of 17 samples exhibited both CAR and PXR agonism (**Table 3**). For the NCI TCM collection, 17 out of 19 samples fell into this category, with all 17 showing both CAR and PXR agonist activity (**Table 4**). Finally, across both CAR modes, 196 out of 300 (65%) and 559 out of 664 (84%) of the DSNP and NCI TCM collections did not meet our initial activity filters, while for both PXR modes, 236 out of 300 (79%) and 504 out of 664 (76%) DSNP and NCI TCM, respectively, did not meet our initial activity filters. Across both the CAR and PXR assays, 154 (51%) and 473 (71%) samples were inactive in all modes for the DSNP and NCI TCM libraries, respectively.

Lastly, we explored the link between our *in vitro* CYP3A4 and PXR assays. Using a subset from both sample collections, a group of 139 CYP3A4 inhibitors made up of our highest quality dose-response curves (potent and efficacious or -1.1 CRC), demonstrated PXR agonism activity, with 100 (72%) also active as PXR agonists. Of the remaining 39, 13 (9%) were CAR agonists, for a total of 113/139 (84%) high quality CYP3A4 inhibitors exhibiting nuclear receptor activity, indicating good agreement between the two assay formats in implicating CYP3A4 (**Figure 3**).

**Figure 3.**
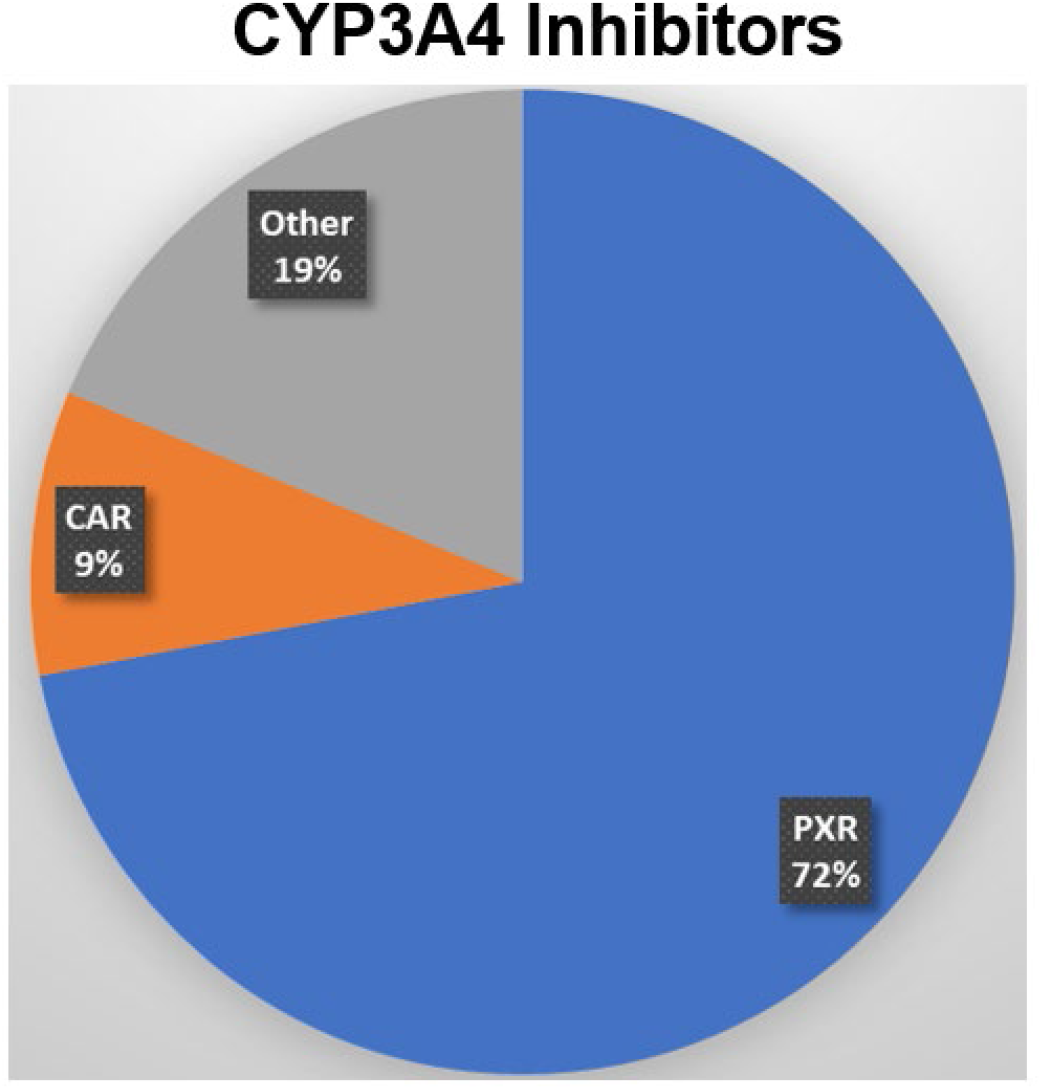
We explored the link between our in vitro CYP3A4 and PXR assays. Using a subset from both sample collections, a group of 139 CYP3A4 inhibitors made up of our highest quality dose-response curves (potent and efficacious or -1.1 CRC), demonstrated PXR agonism activity, with 100 (72%) also active as PXR agonists. Of the remaining 39, 13 (9%) were CAR agonists.

### P-glycoprotein (P-gp) Assay

Together with P450s, transporter-mediated drug interactions are utilized in the characterization of DDIs. We chose P-glycoprotein (P-gp), an ATP-binding cassette (ABC) transporter, due to its importance, prevalence, and potential role in DDIs and/or NaPDI, along with the ability to employ an HTS-amenable assay.^37^ Previous articles have pieced together P-gp information on DNSP and TCM, but none have tested at the scale we describe herein^17-19,38,39^.

To profile both collections for potential P-gp activity, we employed a previously published 1536-well qHTS assay.^40^ Briefly, differential cell viability was measured against two cell lines: a parental KB-3-1 human cervical adenocarcinoma cell line (a HeLa clone) and the drug-resistant subline KB-8-5-11 that overexpresses P-gp. Compounds that kill or inhibit growth of KB-3-1 were expected to demonstrate reduced efficacy against the KB-8-5-11 cells owing to drug efflux by P-gp. The assay performance showed mean Z’-factors of 0.72 and 0.77 for KB-3-1 and KB-8-5-11 cell assays, respectively. The intraplate positive control Bortezomib (a previously identified P-gp substrate^40-42^) performed as expected, with mean IC_50_ values of 16.9 nM (MSR = 1.79) and 62.3 nM (MSR = 1.26) for KB-3-1 and KB-8-5-11 cell assays, respectively, a ∼4-fold difference in potency. Activity was defined as a sample yielding a CRC of -1.1, -1.2, -2.1, -2.2 in the KB-3-1 assay and selective (or potentially interacting with P-gp) if it has a IC_50_ ratio of >5 vs. KB-8-5-11 (if activity was observed).

For DSNP, 19/300 exhibited activity (6%; 15 botanical, 4 supplement; **Table S4a**). Of the 19 active samples, only two substances (<1% tested) exhibited lower efficacy in the KB-8-5-11 cell line (**Figure S5**: the supplement Berberine (a previously identified as a P-gp inhibitor^43^) and botanical Flaxseed Oil with IC_50_’s of 2.0 µM and 1,122 μg/mL, respectively.). While the Flaxseed potency was weak, it does contain polyphenols (lignans), which have been implicated in P-gp modulation.^44^

For the TCM library, 57 out of 664 samples were cytotoxic (9%; 43 organic/14 aqueous; 25 species; **Table S4a**) in the KB-3-1 cell assay. A total of 18 out of 57 samples (covering 8 different species) had the desired outcome of reduced efficacy against the KB-8-5-11 cell line, with an IC_50_ range of 0.2 – 25.1 µg/mL in the KB-3-1 cell assay and weak or inactive vs. the KB-8-5-11 cell line (**Table S4b)**. Most of the selectivity was observed from two species: *Coptis chinensis* (Chinese Goldthread) and *Brucea javanica* (Macassar kernels). All six samples obtained from the *Coptis chinensis* (Chinese Goldthread) species had activity, with IC_50_ value ranges of 0.2 – 2.0 µg/mL and exhibited no activity in the KB-8-5-11 cell line (**Figure S6A**). The other highly active species, *Brucea javanica* (Macassar kernels), had an IC_50_ range of 2.2 – 8.9 µg/mL for its four active samples. *Brucea javanica* samples were weak or inactive in the KB-8-5-11 cell line (**Figure S6B**). A previous publication identified Berberine as a component in *Coptis chinensis*, a possible explanation for its P-gp activity.^45^ For *Bruce javanica*, only the aqueous samples exhibited activity, corroborating earlier work that showed its ability reverse drug resistance in tumor cells.^46^

Results from these studies were compared to previous studies to assess agreement. For example, previous publications identified *Corydalis yanhusuo*, which had two samples with IC_50_ values of 2.2 and 11.2 µg/mL in the KB-3-1 assay (while weak or inactive vs. KB-8-5-11), as an inhibitor^47^, but no difference in cell viability was observed in the other four samples found in the collection. *Arctium lappa* (Greater burdock) had two samples from an aqueous fraction exhibit apparent P-gp activity with IC_50_ values 8.9 and 13.7 µg/mL (weak or inactive vs. KB-8-5-11), but no difference in cell viability was observed in the other four samples found in the collection.^48^ Single sample organic extracts of *Trachelospermum jasminoides* (star jasmine; contains the bioactive compound Astragalin^49^) were detected in our KB-3-1 screen with an IC_50_ value of 6.3 µg/mL (weak activity in KB-8-5-11 assay), and similar cell viability was observed in the remaining five samples. Single samples from the organic extracts of *Aster tataricus* (Tatarian aster), *Melia toosendan*, and *Menispermum dauricum* (Asian moonseed) exhibited IC_50_ values 25.1, 13.5, and 20.3 µg/mL (weak or inactive vs. KB-8-5-11), respectively, but no difference in cell viability was observed in the remaining samples.^50^ *Menispermum dauricum* contains dauricine which has been identified as a P-gp modulator. Further investigation into these singular extract solutions might identify the component(s) responsible for the difference in cell viability.

### Viability Assay using HepG2 Cell Line

To assess broad cytotoxicity, we employed screening against a HepG2 cell line, as previously described.^51^ Briefly, cell viability over a 24-hour time-period was determined using both GF-AFC substrate and Cell-Titer Glo (CTG), measuring cellular protease activity and ATP content, respectively. This is similar to the incubation time and readout (GF-AFC) chosen for the CAR and PXR assays, with the major difference the concentration range tested; CAR and PXR assays were tested up to 90 μM, while the HepG2 assay was tested up to 30 μM (or 0.2 mg/mL and 4 mg/mL for powder mixtures and oils, respectively). The assay performance was acceptable, with mean Z’-factors of 0.53 and 0.77 for GF-AFC and CTG cell assays, respectively. In the GF-AFC assay, a lone DSNP sample exhibited activity, lithospermic acid (a constituent of Comfrey Root, NMIX00599553; inactive), with an EC_50_ of 0.8 μM. For the NCI TCM collection, 21 out of 664 samples (3%) exhibited activity in the GF-AFC assay, ranging in potency from 0.05 to 35.8 μg/mL. In the CTG assay, no DSNP samples were cytotoxic, while 53 out of 664 TCM samples (8%) were identified as toxic, ranging in potency from 0.28 to 38.1 μg/mL (**Table S5**).

To determine if a sample was cytotoxic, we looked for overlap between the two assays, identifying 10 samples, all from the NCI TCM as cytotoxic. The 10 samples were comprised of seven organic and three aqueous extracts from nine species, with *Vaccaria segetalis* (Wang Bu Liu Xin or Cowherb seed) being the only species having >1 sample appear, with EC_50_ values of the two aqueous samples ranging from 15.8 – 21.4 μg/mL. Based on our criteria, the DNSP library exhibited no cytotoxicity over the 24-hour time-period in HepG2 cells. The NCI TCM collection was largely inactive, with only 1.5% of samples exhibiting cytotoxicity.

### Clustering and PCA Comparison vs. NCATS Pharmaceutical Collection (NPC)

Compounds that show similar bioactivity profiles tend to share similar targets or mechanisms-of-action (MOA).^25,52,53^ The Toxicology in the 21st Century (Tox21) group at NCATS performed a similar panel of *in vitro* assays^52^ on the NCATS Pharmaceutical Collection (NPC), a comprehensive collection of clinically approved drugs.^54^ To compare the activity profiles of the DSNP and NCI TCM with those of the NPC across the same panel of assays, we clustered the NPC, DSNP, and NCI TCM samples based on similarity in their bioactivity profiles (measured by curve rank, **Table S6**), using the CYP450, CAR, PXR, and P-gp assays **(Figure 4)**. In addition, we performed a PCA on these activity profiles to visualize DSNP and TCM samples in comparison to NPC across the various endpoints (**Figure 5**). Both the PCA and clustering analysis results show that these three collections largely overlap many samples that share similar activity profiles, thus suggesting similar MOAs. These data display co-clustered activity profiles among subsets of samples tested, represented by clustering numbers (**Table S6**). We then identified the prominent MOAs enriched in each cluster. The statistical significance of enrichment is measured by the p-value from the Fisher’s exact test. Overall, we saw 30 significant MOAs in 17 bioactivity profiles, containing 78, 284, and 745 samples from DSNP, TCM, and NPC, respectively (**Table 5**). Many of the drugs in the NPC have well-annotated MOAs, while the MOAs for most of the DSNP and TCM samples remain unknown. Samples with similar bioactivity profiles to well characterized drugs are likely to share similar biochemical interactions, therefore we can hypothesize the MOAs of the DSNP/TCM samples as they may share the same MOAs as their co-clustered NPC drugs.

**Figure 4.**
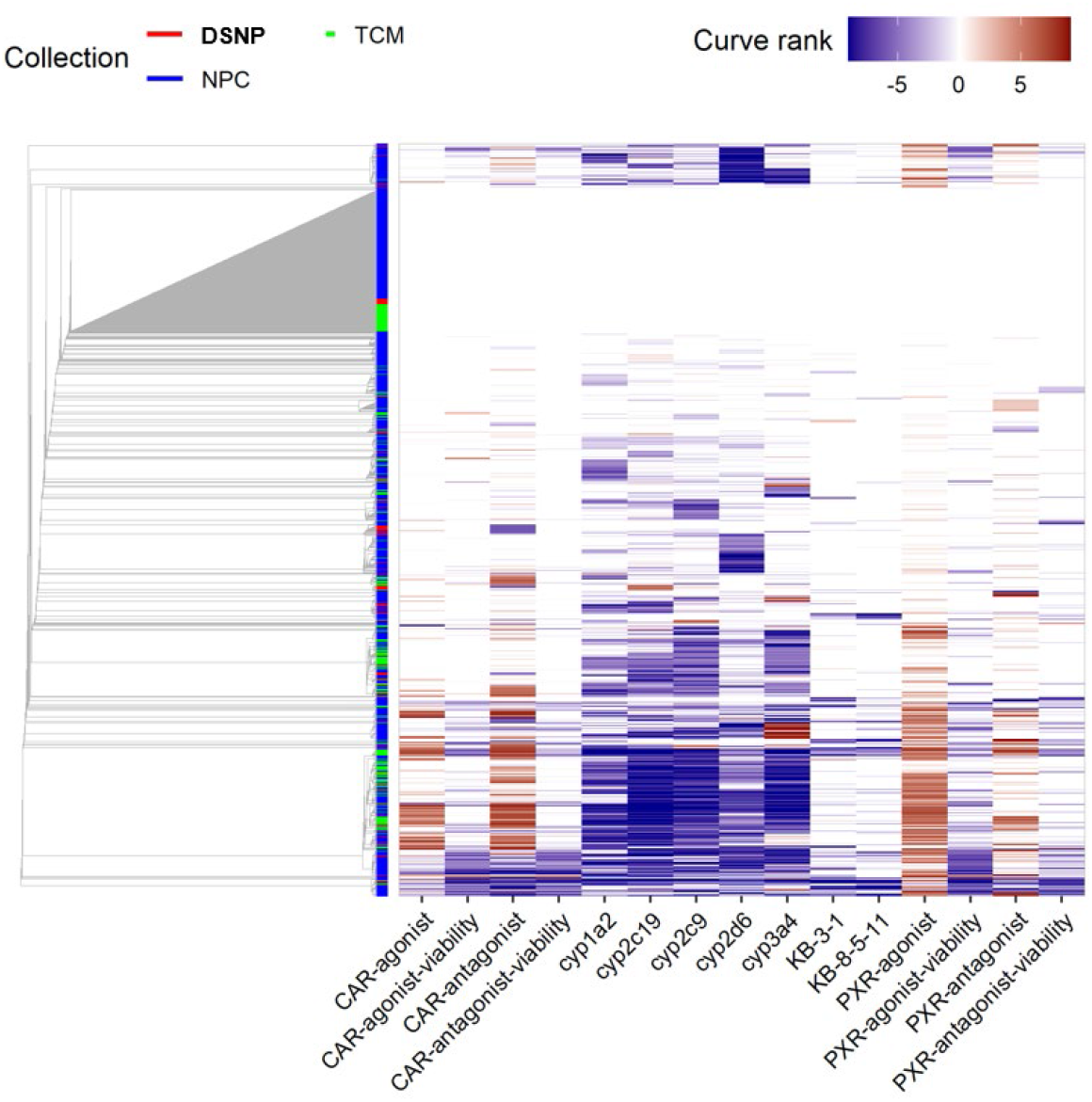
Clustering based on activity profile similarity: The DSNP and TCM compounds tend to co-cluster with the NPC compounds.

**Figure 5.**
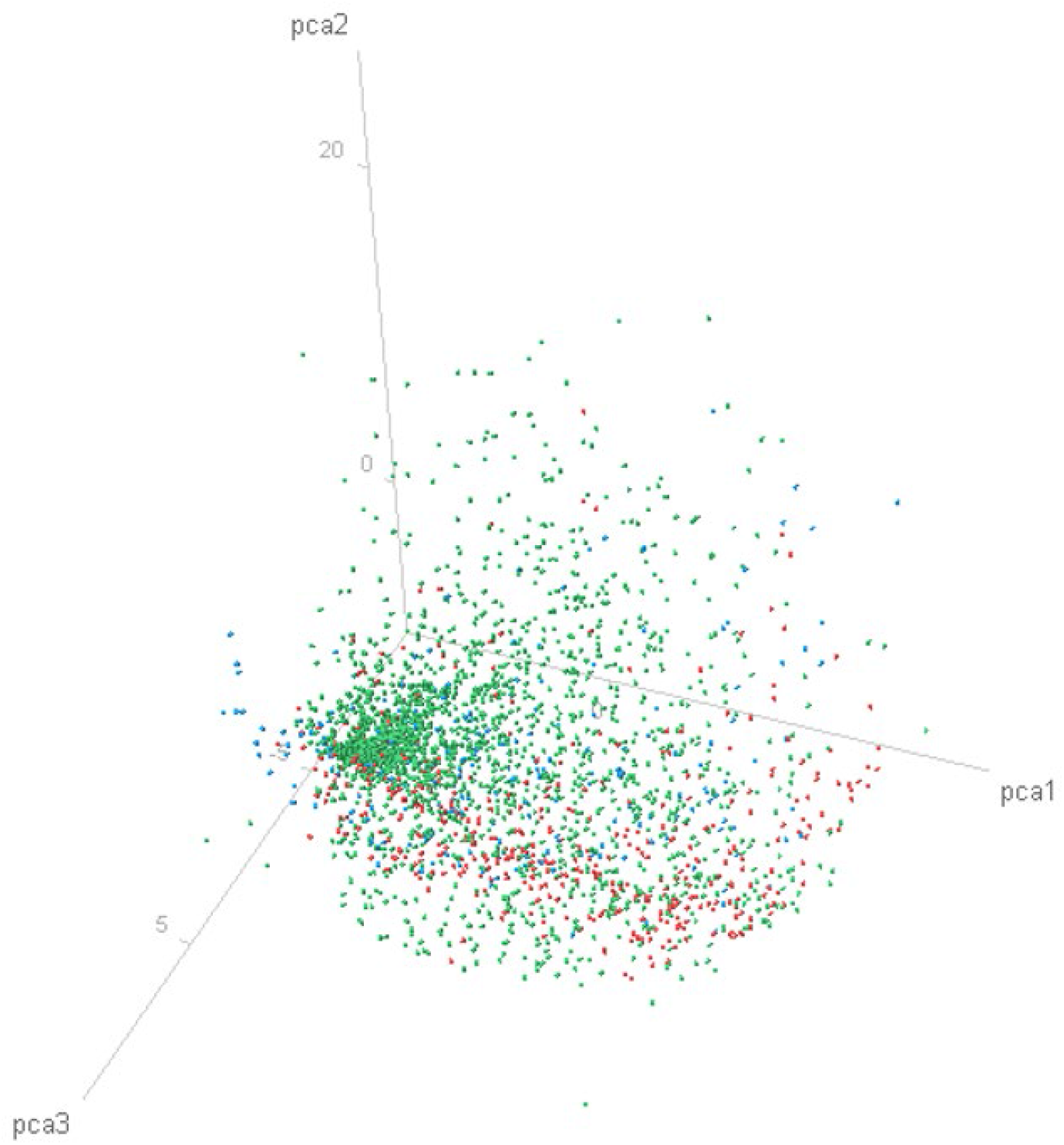
The activity spaces covered by DSNP 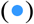 and TCM 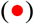 fall within that of NPC 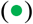.

We chose to highlight four bioactivity profiles based on a measured p-value < 1 × 10^−6^ and having >10 samples, as examples of this prediction. As a first example, bioactivity profile #482 contained five substances each from the DSNP and TCM collections, associated by their strong inhibition of CYP2D6 and lack of activity in the other assays. Two of the highest p-values co-clusters were histamine receptor antagonists and adrenergic receptor agonists, with p-values of 1.32 × 10^−7^ and 7.52 × 10^−6^, respectively. The MOA of these related to treatments for allergy or GI issues (with side effects like somnolence) and medications that that relax muscles of the airways, causing widening of the airways resulting in easier breathing, respectively. Three of the five DSNP samples are widely used in the US population: the supplements Choline Chloride (NCGC00095059), alpha-Tochopherol (NCGC00142625), and Doconexent (Docosahexaenoic acid, DHA; NCGC00161345). While the exact MOA of these supplements is unknown, choline chloride might play a role in cardiovascular and neurological disorders, alpha-Tocopherol acts as an antioxidant that has been linked to a litany of diseases, and DHA is an omega-3 fatty acid believed to support healthy brain development in young children, prevent cardiovascular disease and cognitive decline during Alzheimer’s disease. *Corydalis yanhusuo* (three aqueous) and *Stephania tetrandra* (two aqueous) from the TCM library were included in the cluster, having been used for analgesic, sedative, hypnotic, headache, insomnia, dysmenorrhea, and arthralgia caused by rheumatism, wet beriberi, dysuria, eczema and inflamed sores, respectively.

A second bioactivity profile #414, containing 29 DSNP and 127 TCM samples, co-clustered with sterol demethylase inhibitor (antifungals) and insecticides (i.e. Allethrin) with p-values of 1.81 × 10^−5^ and 5.18 × 10^−5^, respectively. The bioactivity characteristics of this cluster were active in all CAR and PXR modes, along with pan-CYP activity. This was the largest co-cluster, representing ∼10% of DNSP and ∼19% of TCM samples. Notable from the DSNP Library is Saw Palmetto (NMIX00599537), a widely used botanical for urinary symptoms associated with an enlarged prostate gland as well as for chronic pelvic pain, migraine, and hair loss.^55,56^ Notable among the TCM samples were *Lindera aggregata* (three organic and one aqueous) and *Chrysanthemum morifolium* (four organic). *Lindera aggregata* has been used to treat chest and abdomen pain, indigestion, regurgitation, cold hernia and frequent urination, while *Chrysanthemum morifolium* has been used in treatment of inflammation, hypertension, and respiratory diseases.

A third bioactivity profile #456 exhibited pan-CYP activity along with unwanted cytotoxicity in the nuclear receptor assays. This cluster contains nine DSNP and six TCM samples, co-clustered with anti-infective agents (local; inhibit the activity of harmful microorganisms such as Cetylpyridinium chloride and Chlorhexidine dihydrochloride) and topoisomerase inhibitor (therapeutics against infectious and cancerous cells), with p-values of 1.83 × 10^−5^ and 3.12 × 10^−5^, respectively. Notable from the DNSP library are the botanical Ashwagandha (NMIX00599593) and supplement Omega-3 fatty acids (NMIX00599569). Ashwagandha is best known for its alleged ability to reduce stress, and while Omega-3 fatty acids have proven cardiovascular benefits, this would suggest the latter, at high concentrations would need to be studies further in the CAR and PXR assays. This effect did not appear for DHA (NCGC00161345), indicating the mixture appears to have negative synergy at high concentrations. For TCM, two organic samples each from *Dioscorea nipponica* and *Rubia cordifolia. Dioscorea nipponica* has been used to prevent and treat coronary heart disease, while *Rubia cordifolia* medicinal uses include treatment of skin diseases like eczema, dermatitis and skin ulcers. The latter, as a local agent, might have a common MOA.

Finally, a fourth bioactivity profile #476 (CYP2C9 inhibition), containing one DSNP and nine TCM compounds, co-clustered with cyclooxygenase inhibitors (non-steroidal anti-inflammatory drug or NSAID), with a p-value of 5.35 × 10^−5^, highlighted by Peanut Oil (NMIX00601620) and two aqueous samples from *Artemisia argyi*. This indicates peanut oil, typically used in cooking, could have anti-inflammatory effects. *Artemisia argyi* is mainly used in TCM for treatment of “irregular menstruation, infertility, hematemesis, epistaxis, metrorrhagia and pruritus”.

In summary, exploiting the predicted MOAs based on the present bioactivity profiles could lead to more focused research and potential avenues to understand the biological activity patterns of herbs and mixtures.

## Discussion

Dietary supplements have been woven into human culture as potential treatments for a variety of ailments. Despite their remarkable high level of use, their bioactivity characterization compared to approved drugs is minimal. For example, a paradox exists: while the drug approval process has largely been a highly detailed characterization of a single component and its interaction pharmacokinetically and dynamically with the human body, the supplement and herbal process has largely been one of anecdotal use and little rigorous scientific research.^57,58^ We addressed this challenge by designing and assembling a collection of DSNP along with the NCI TCM collection and employing these collections into screens using assays of broad relevance to substance toxicity and metabolism. qHTS analysis established a rich set of biological profiles and this data allows us to further assess toxicity and explore potential MOAs. This process translates these basic findings into information that can eventually be used by clinicians to better provide patients with accurate information on the efficacy and safety of supplements.

The ability for prospective researchers to procure the same substance tested is of extreme importance in the field of supplement and botanical research, due to historical difficulties of researchers attempting to repeat previous works on these types of substances and their varying degree of availability and integrity. Organizations have attempted to address this problem; for example, the Dietary Supplement Analytical Methods and Reference Materials Program (AMRM) has developed over 55 reference materials for researchers to procure since its inception.^59^ We expanded on these efforts by incorporating commercial vendors in our Dietary Supplements and Natural Products (DSNP) library. By identifying commercial vendors for the substances presented in this work, we hope to accelerate the procurement process for the research community to enable better reproducibility and subsequently improved characterization of their biological activity outside of this work. Shortly after we assembled the DSNP library, the NaPDI Center published a recommended list of 47 natural product candidates that should be studied.^60^ Serendipitously, our group used a similar source for identifying natural products as NaPDI (*HerbalGram*, Journal of the American Botanical Council^15^, University of Washington Drug Interaction Database).^61^ Overall, we feel the substances included in our DSNP library appear to be ones of most interest with flexibility to add and/or subtract for future testing.

There are a multitude of difficulties and challenges faced by clinicians when presented with patients who are taking Chinese medicines in combination with an approved drug or OTC medication.^62^ While there are a small number of examples from literature of plant extracts that contain useful clinical data, given the thousands of extracts available, more characterization needs to be done in this field. Adding to the difficulty is that these extracts can be prescribed as a mixture with other extracts of unknown bioactivity, compounding the uncertainty of their efficacy. NCI was able to obtain 132 different species in putting together the TCM collection, totaling 664 extract samples, and was assembled with great care. Chinese medicine also includes the use of formulas and decoctions, with the composition containing multiple plant extracts and changing properties based on the preparation of the mixture.^63,64,65^ Given the influence eastern medicine has on western culture, we felt it was important to include TCMs in our panel of assays as they should be characterized in a similar manner to an approved drug, with one trying to determine not only if there is a therapeutic efficacy of these samples, but what the therapeutic window is when compared to any liabilities.^66^

We first employed our screening capabilities against the cytochrome P450 enzyme family, which performs essential functions for normal metabolism, influences drug pharmacokinetics, and affects negative outcomes in patients through adverse drug-drug interactions. This enzyme family is responsible for the metabolism of approximately two-thirds of known drugs in humans, with 74% of this attributable to five isozymes: CYP1A2, CYP2C9, CYP2C19, CYP2D6, and CYP3A4.^67^ We chose the HTS-amenable P450-Glo kit, while noting that an inhibitory outcome could result from a compound acting as either an inhibitor or substrate because both will compete for an enzyme.^21^

Due to the different assembly/construction of the two libraries, we performed analysis separately for each. The DSNP library exhibited activity across all CYPs tested, with 2C19 > 1A2 > 2C9 > 3A4 > 2D6. Overall, 64% of the DSNP samples tested had a role in one of the major human P450 metabolizing enzymes with CYP2D6 being the least likely. Given their wide prevalence of historical use, this suggests that the human liver evolved to metabolize and clear these substances from the body through a combination of multiple CYPs.

Examining the relationship between constituents and their parent mixtures, the two examples we highlight contained the largest number of constituents (18) associated with a mixture, from Saw Palmetto and Peony Root Powder. For the pan-active Saw Palmetto, three compounds (Linoleic acid, Oleic acid, Lauric acid) were possibly responsible for a majority of its response, except none were active against CYP2D6. Our *in vitro* biochemical screening data differs from a previous serum study, where no significant effect on CYP1A2, CYP2D6, CYP2E1, or CYP3A4 activity was observed.^56^ The parent substance Peony Root was inactive while its components, Pentagalloyl glucose and NP-003686, exhibited strong pan-activity. While we spent our efforts examining for synergistic effects among the constituents, Peony Root might represent an example of anti-synergistic effect, where the mixture of components interacts and negates each other. These examples highlight the difficulty in translating the constituent concentrations (known) to their respective mixtures (typically not known), and it validates the ongoing research in the natural products field to identify purified components and determine their influence on the bioactivity of mixtures.^68^ A recent example of this is the supplement Cannabidiol (CBD) which was pan-active in our *in vitro* P450 assays. This supplement was part of a group of Cannabinoid acids isolated from hemp (*Cannabis sativa*) that were tested and successfully blocked the cellular entry of SARS-CoV-2.^69^

Chinese herbs are traditionally taken as an aqueous extract (i.e., decocted as tea and also given as tincture [1:3 to 1:5]).^70,71^ Since aqueous extraction is typically used, our *in vitro* findings suggest minimal NaPDI since 67% of the aqueous extracts displayed no activity over the concentration range tested. This left a subgroup of 19 species having all aqueous samples exhibit inhibitory activity against any CYP isozyme. Only one species, *Lindera aggregata*, exhibited pan-activity, and as such is the leading candidate mixture to potentially have an NaPDI, since pan-activity would be an undesirable characteristic due to potential NaDPIs. The only other species to have strong aqueous extract activity was *Rosa rugosa*, a plant with antimicrobial properties that is used to treat a wide variety of ailments such as chronic inflammatory disease.^72^ We surmise that the aqueous extraction became the favored method due to less toxicity, as we observed 5- to 7-fold higher activity across all CYPs tested. The organic extracts exhibited activity across all CYPs tested, as 1A2, 2C9, 2C19, and 3A4, had similar number of inhibitors identified, ranging from 85% – 91%, with 2D6 exhibiting the lowest number of inhibitors at a still impactful 52% of mixtures received. In contrast, these numbers sharply decreased with the aqueous extract activity, ranging from 10% - 19%. Overall, these numbers indicate that the organic extracts yielded more of the components that modulate CYP activity, reinforcing the importance of how these medicines are prepared and how it can impact their resulting bioactivity and/or how future extraction work to identify active ingredients should be performed.^73^

We further characterized both libraries in CAR and PXR cellular assays which regulate the expression of several CYP450 enzymes. Activation or inhibition of CAR and PXR has been demonstrated to cause modulation of CYP enzymes as well as play major roles in energy homeostasis, cancer, and immune response.^74^ Identifying natural and therapeutic compounds as modulators of these important nuclear receptors can significantly reduce and potentially eliminate DDI and NaPDI, as well as improve our understanding of their potential therapeutic benefits.

Comparing these cellular experiments to the biochemical CYP3A4 assay, 84% of our highest quality CRCs from the biochemical screen exhibited nuclear receptor activity, indicating good agreement between the two assay formats. In the DSNP collection, St Johns Wort, one of the more well characterized botanicals identified as a CYP3A inducer^75,76^, was identified as a PXR agonist (while also exhibiting weak CAR agonism), along as a CYP1A2, CYP2C9, CYP2C19, and CYP3A4 inhibitor. A similar bioactivity profile was observed with Resveratrol, a previously researched P450 modulator.^25,77^ Resveratrol has been claimed to help with life extension and is responsible for some of the health benefits associated with red wine. Both Kava Kava Root and Holy Basil exhibited dual nuclear receptor agonist activity along with activity in all the P450 inhibitory assays. While Kava Kava root has been previously characterized in a high-throughput setting^25^, there is very little known on the CAR and PXR activity of Holy Basil, which has been suggested as an anti-inflammatory. The ability to link constituent activity to parent activity was not as clear cut in these assays, with only one of three highlighted having a concordant result.

For the NCI TCM collection, we highlighted a single CAR antagonist derived from an aqueous extract of *Polygala tenuifolia*, which has traditionally been used for its protective effects on the brain. In our assay, the extract was inactive against all CYP450 tested.^78^ Approximately 16% of the collection (comprising 50 species) exhibited CAR agonism, with the most potent an organic extract of *Cynanchum atratum* which has been used in folk medicine for skin inflammatory diseases.^79^ This extract exhibited inhibitory activity with CYP1A2, CYP2D6, and CYP3A4. Similarly, 24% of the collection yielded a PXR agonist response, with two species from organic extracts exhibiting the most activity. *Ligusticum chuanxiong* and *Evodia rutaecarpa* (Fruits of Evodia, Wu Zhu Yu) were both pan-active in the *in vitro* P450 assays. *Ligusticum chuanxiong* was previously identified as a PXR activator and is traditionally used in Chinese medicine for the treatment of cardiovascular diseases, headache, and vertigo.^80^ *Evodia rutaecarpa*, which has been recommended to increase body temperature^81^, has been show to activate human CAR due to the alkaloid evodiamine and metabolize mouse CYP1A2.^82 83^ While these mixtures exhibit a considerable amount of bioactivity, their promiscuous nature makes it difficult to determine which component of the extract contribute to nuclear receptor activity. Nonetheless, the bioactivity profiles of these extracts may serve as a starting point for a probe or drug discovery campaign. Additionally, compounds/mixtures can inhibit CYPs directly (not Glo-assay inhibition) by competitive or non-competitive inhibition. Compounds that inhibit CYP via competitive/noncompetitive inhibition as well as activate/inhibit nuclear receptors (one or both) that regulate these CYPs would most likely have high DDI-potential and complex outcomes in the clinical needs.

We then pivoted to the drug transporter P-gp due to its importance, prevalence, and potential role in DDIs.^37^ Only two substances from the DSNP library (<1% tested) met the criteria for inhibition, the supplement Berberine (a previously described P-gp inhibitor) and the botanical Flaxseed Oil. Berberine, a substance found in goldenseal, has been studied for heart failure, diarrhea, infections, and other health conditions. The activity from Flaxseed Oil, which may assist in cardiovascular disease and type II diabetes, might be due to polyphenols which have been known to influence P-gp.^84,85^ For the TCM samples, the majority of the selectivity was observed from two species: *Coptis chinensis* (Chinese Goldthread) and *Brucea javanica* (Macassar kernels). A previous publication identified Berberine as a component in *Coptis chinensis*, a possible explanation for its P-gp activity, where the rhizome (or huanglian) is alleged to help with multiple illneses.^45,86^ For *Bruce javanica*, due to the aqueous samples contributing to the activity, we would recommend further exploration on its potential for P-gp related toxicity, even though this plant was earlier described in the “treatment of lung, prostate, and gastrointestinal cancer, and has potent antimalarial, anti-inflammatory, and antiviral effects, with low toxicity”.^46^ Due to screening limitations, only cytotoxic or cytostatic P-gp substrates were identified, which resulted in few samples exhibiting activity in this assay. To mitigate these limitations, our future efforts will incorporate our acoustic dispensing matrix platform, enabling us to add known substrates or inhibitors (i.e., Tariquidar) in dose-response along with library samples plated in dose-response to observe possible shifts in potency. As previously noted, the need for compounds to demonstrate cytotoxicity is a limiting factor, so by having them displace a known inhibitor one could potentially identify more P-gp modulators.

The above panel of assays were utilized to represent an efficient means to interrogate NaPDI/metabolism from prominent human metabolism pathways to assist in prioritizing and evaluating the substances in the DSNP and TCM collections for further evaluation. Using the bioactivity results from each assay, we performed PCA and cluster analysis to generate a bioactivity profile for each substance. These profiles were then combined with previous screening results against our approved drugs collection to predict potential mechanism-of-action (MOA), totaling 30 enriched MOAs, with the glucocorticoid receptor agonist and serotonin receptor antagonist both appearing in three clusters. The rich set of biological profiles established in this study allows researchers to further evaluate potential toxicity of substances in the library. For example, peanut oil was clustered with NSAIDs and showed promise in an investigation for inflammatory properties in mice with type II diabetes because peanut oils are high in oleic acid. Collectively, these findings could lead researchers to prioritize anti-inflammatory models as a next step in determining peanut oil activity *in vivo* and further make recommendations to continue or terminate use. A more complicated consideration arises from the plant extract *Lindera aggregata*, which was co-clustered with sterol demethylase inhibitor (antifungals) and insecticides. Earlier scientific work tested the root tubers of *Lindera aggregata* and identified compounds with insecticidal activity.^87^ While a notable find from our analysis, extracts from this plant have been claimed for cardiovascular, cancer, and diabetic treatments, along with anti-inflammatory effects of components that were isolated, purified, and charaterized^88^; somewhat concerning is the relations to insecticides, a class of substances that is not recommended for use in humans. Results derived from this work suggest that a portion of the DSNP and TCM substances tested exhibit some degree of activity, and our understanding of these substances would benefit from additional research.

In summary, we assembled a collection of commercially available DSNP and with the existing TCM collection characterized them against a panel of *in vitro* assays (CYP450, CAR, PXR, and P-gp assays) to assess their effects on drug metabolism and transport pathways. The activity profiles of these substances against the CYP isoforms, which play important roles in drug metabolism, revealed that constituents from the DSNP library and organic extracts from the TCM library exhibited the most potent activity against any one isozyme. Moreover, substances that showed pan-activity against all five CYP isozymes were found to be two-folds higher in the TCM library compared to the DSNP library. We examined the activities of constituents and their parent mixtures and found that, in some instances, constituents showed activities different from their parents. Activity profiles of substances in the DSNP and TCM collections were compared against well-characterized drugs in the NPC through cluster analysis and PCA. Many substances in the three collections showed similar activity profiles, suggesting shared MOAs. The data collectively accrued in this study can assist in the prioritization and evaluation of samples in the DSNP and TCM collections for existing and future efforts aimed to better predict safety and efficacy profiles of supplements and herbal medicines.

## Acknowledgements

We thank NCI, Beijing University of Chinese Medicine, and Harvard as the source of the material and that collection of material was partially supported by U19CA128534 from NCI in any publication of research findings resulting from the use of the material. We would like to thank the following colleagues for their support and input: Matthew D. Hall, Jim Inglese, Ken Cheng, Jason Rohde, Noel Southall, Xin Xu, Carina Danchak, Dac-Trung Nguyen; from NCI Carol Haggerty and Barry R. O’Keefe.

## Materials and Methods

### Materials

Assay reagent components were procured from Promega, Sigma, or ThermoFisher unless otherwise specified.

### Library Assembly

Selected substances were cross-referenced with information primarily from NCCIH^89^, National Library of Medicine (NLM)^90^, Operation Supplement Safety (OPSS)^91^, and Spectrum Chemical Botanical Dietary Ingredients^92^, along with reporting from new agencies.^93-96^ This was done to synchronize nomenclature and reference information among the multiple sources, where we initially identified 301 substances (**Table S7a**). We then applied additional criteria such as fitness for high-throughput screening (HTS; solubility, solution viscosity), commercial availability, and cost, leading to 131 substances that were procured, followed by categorization as either botanicals (88 samples) or supplements (43 samples) as defined by the NIH Office of Dietary Supplements (ODS^97^; **Table S7a**). Finally, the GSRS database^98^ was used to associate those substances with reported purified components (“constituents”; **Table S7b**) to see if any of the purified components can be linked back to the activity of a substance mixture. We sourced an additional 169 compounds (for a total of 300 samples **Table S7c**) representing 39/131 substances (either botanical or supplement) that had components identified.

### Dietary Supplement and Natural Product (DSNP) Library Preparation

Dietary Supplements (vitamins and minerals), and Natural Products (herbal and plant extracts; botanicals) were commercially sourced by NCATS. Substances were processed by the NCATS Compound Management group. Briefly, powders were manually weighed in 20 mL amber glass vials, solubilized with DMSO, then capped and vortexed for 10 seconds at 3,200 rpm. Solutions were visually inspected to ensure the mixture completely dissolved; if not, solution was gently warmed to 50°C and sonicated for up to 10s in a water bath to assist in dissolution. Contents were then transferred to a new Matrix 2D barcoded sample tube, securely capped, and placed in the Automated Compound Store.

Solubilization of the substances was achieved using the methodology described in Butler *et al*. “Natural Product Libraries: Assembly, Maintenance, and Screening” from 2014.^99^ For dry powders, we chose a top concentration of 50 mg/mL as, according to their methodology, this would represent an approximate final assay concentration of 50 µM, the standard top concentration our Center performs HTS assays. All oils were assumed to be a density of 1 g/mL unless otherwise supplied by the vendor, corresponding to an approximate final assay concentration of 1,000 µM. DMSO was chosen as the solvent for solubilization along with compatibility with the pin types used on our Wako Pin-Tool instrument, and our Center’s long-standing practice of storing compounds for screening in DMSO, and compatibility with our acoustic liquid handlers.

The library was formatted into 1536-well plates in a modifed-qHTS format, where each sample underwent interplate serial dilutions (1:2, 12-points) in 384-well plates, for a final source concentration ranging from 24.4 µg/mL to 50.0 mg/mL and 488 μg/mL to 1,000 mg/mL for powders and oils, respectively. The samples were then stamped into different quadrants of three 1536-well plates, with the first plate containing the top four concentrations, the second plate the next four concentrations, and the third plate the last four concentrations. Plates were stored in a desiccator at room temperature protected from light.

### NCI Library of Traditional Chinese Medicinal (TCM) Plant Extracts Preparation

The NCI Library of Traditional Chinese Medicine (TCM) Plant Extracts was kindly provided by the Natural Products Branch of the Developmental Therapeutics Program in the Division of Cancer Treatment and Diagnosis at the NCI. The library contained 332 extracts representing 132 species, extracted from aqueous and organic solvents, totaling 664 samples, and was provided as eight 96-well plates (four aqueous, four organic; **Table S7d**). Each sample arrived as a 500 μg film and were solubilized with 33 µL of DMSO for a top concentration of 15 mg/mL, as previously performed.^63^

The library was formatted into 1536-well plates in a modifed-qHTS format, where each sample underwent interplate serial dilutions (1:3, 12-points) on 384-well plates, totaling 12 concentrations for a final source concentration range of 0.0847 µg/mL to 15.0 mg/mL. The samples were then stamped into different quadrants of three 1536-well plates, with the first plate containing the top four concentrations, the second plate the next four concentrations, and the third plate the last four concentrations. Plates were stored in a desiccator at room temperature protected from light. Samples are sent anonymized; upon identification of an active, NCI Natural Products Branch provided taxonomy and type of extract.

### P450 assays

Assay conditions were adopted from previously described methods incorporating the P450-Glo™ Assay (Promega, Madison, WI).^23,100-104^ Briefly, a 2 µL mixture of enzyme and luciferin-substrate (final concentrations CYP enzyme-specific and pro-luciferin substrate-specific) or minus-enzyme (assay buffer conditions supplied by manufacturer for respective enzyme isoform; typically 100 mM KPO_4_) and luciferin-substrate were dispensed using a BioRAPTR FRD™ (Beckman Coulter, Brea, CA) into columns of a 1536-well white solid-bottom medium binding plate (Greiner Bio-One, Monroe, NC). Substances (final concentrations of most substances ranged from 19.5 nM – 39.8 μM) and controls (CYP-specific) were transferred (16 nL) via pintool (Wako Automation, Richmond, VA) and incubated (RT) for ∼15 minutes. Next, 2 μL of NADPH regeneration solution (final concentration 1X) was dispensed to initiate the reaction, which then proceeded for 30 – 45 minutes (RT, protected from light), allowing NADP^+^ to convert to NADPH, which then serves as the electron source for the CYP oxidative reactions leading to the pro-luciferin substrate to convert to D-luciferin.^104^ Next, 4 µL of reconstituted Luciferin Detection Reagent was dispensed to allow for D-luciferin to be converted to light via luciferase reaction. Samples were centrifuged for 15 seconds at 1,000 RPM’s to remove bubbles, followed by a RT incubation for 30 minutes. Plates were then read for luminescence intensity on a ViewLux detector (Perkin Elmer; Waltham, MA).

To account for detection kit artifacts, compounds were counterscreened for interferences using previously described methods.^105-108^ Briefly, 4 µL of D-Luciferin (final concentration 1 µM) or buffer (100 mM Potassium phosphate) were dispensed into a 1536-well white solid-bottom medium binding plate. Compounds (final concentration of 19.5 nM – 39.8 μM) and intraplate control PTC124 (final concentration range 1.2 nM – 39.8 µM) were transferred (16 nL) via pintool (Wako Automation), for a and incubated (RT) for ∼15 minutes. Next, 4 µL of reconstituted Luciferin Detection Reagent was dispensed, samples were centrifuged for 15 seconds at 1,000 RPM’s and incubated (RT) for 30 minutes. Plates were then read for luminescence intensity on a ViewLux detector.

### CAR and PXR reporter gene assays

Assay conditions were adopted from previously described methods.^26,109^ For CAR luciferase reporter gene assays, HepG2-CYP2B6-hCAR cells were cultured in DMEM (Invitrogen) supplemented with 10% Hyclone™ FBS (GE Healthcare Life Sciences, Logan, Utah), 5 µg/ml blasticidin (Invitrogen, Carlsbad, California), 0.5 mg/ml geneticin (Invitrogen), and 100 U/ml penicillin and 100 μg/ml streptomycin (Invitrogen). Briefly, 4 µL of HepG2-CYP2B6-hCAR cells at 6.25 × 10^5^ cells/mL (2,500 cells/well) were dispensed using a Multidrop Combi dispenser in tissue culture–treated 1536-well white assay plates (Greiner Bio-One) in the same culture media, without geneticin and blasticidin included. Plates were incubated for 5 hours at 37°C, 95% humidity, and 5% CO_2_ to allow for cell attachment. Compounds (final concentrations of most substances ranged from 15.6 nM to 31.9 μM), positive control (for agonist mode: 6-(4-Chlorophenyl)imidazo[2,1-b][1,3]thiazole-5-carbaldehyde O-(3,4-dichlorobenzyl)oxime (CITCO) with a final concentration range 2.8 nM to 92 μM; for antagonist mode: PK11195 with a final concentration range 2.8 nM to 92 µM), and toxicity control tetraoctyl ammonium bromide (final concentration of 92 µM) were transferred (23 nL) via pintool (Wako Automation). After compound transfer, plates were dispensed with 1 µL of the known hCAR antagonist PK11195 (final concentration of 0.75 µM) in the agonist mode, or 1 µL of CITCO (final concentration of 48 μM) in the antagonist mode, using a BioRAPTR FRD™ and incubated for ∼24 hours at 37°C and 5% CO_2_. Following incubation, 1 µL of CellTiter-Fluor™ (Promega, Madison, Wisconsin) was added using a BioRAPTR FRD™, after which, all plates were incubated (37°C/5% CO_2_) for 1 hour then measured for fluorescence intensity (E_x_/E_m_ = 405 nm/540 nm) on a ViewLux plate reader to determine cell viability. Directly after this fluorescence reading, 4 µL of ONE-Glo™ Luciferase reagent (Promega) was added to each well using the BioRAPTR FRD™, followed by a 30-minute incubation (RT) and read for luminescence intensity on a ViewLux detector.

For PXR luciferase reporter gene assays, hPXR-Luc HepG2 cells provided by Dr. Taosheng Chen (Department of Chemical Biology and Therapeutics, St. Jude Children’s Research Hospital) were used.^110^ Cells were cultured in EMEM supplemented with 10% fetal bovine serum, 100 U/ml penicillin and 100 μg/ml streptomycin, and 500 μg/ml of geneticin. Briefly, 5 µL of hPXR-Luc HepG2 cells in phenol red-free DMEM containing 5% charcoal stripped FBS, 1 mM sodium pyruvate (Invitrogen), 2 mM L-glutamine (Invitrogen), and 100 U/ml penicillin and 100 μg/ml streptomycin were dispensed using a BioRAPTR FRD™ at 4 × 10^5^ cells/mL (2,000 cells/well) in tissue culture–treated 1536-well white assay plates (Greiner Bio-One). Plates were incubated for ∼5 hours at 37°C, 95% humidity, and 5% CO_2_ to allow for cell attachment. Compounds (final concentration most substances ranged from 15.6 nM – 45.9 μM), positive control rifampicin (RIF; final concentration range of 2.8 nM – 92 µM) for the agonist mode or SPA70 (final concentration range of 2.8 nM – 92 µM) for the antagonist mode, and cytotoxicity control tetraoctyl ammonium bromide (final concentration of 92 µM) were transferred (23 nL) via pintool (Wako Automation). After compound transfer, plates were dispensed with 1 µL of the media in the agonist mode, or 1 µL of RIF (final concentration of 2 µM) in the antagonist mode, using a BioRAPTR FRD™ and incubated for ∼24 hours at 37°C and 5% CO_2_. Following incubation, 1 µL of CellTiter-Fluor™ (Promega, Madison, Wisconsin) was added using a BioRAPTR FRD™, after which, all plates were incubated (37°C/5% CO_2_) for 1 hour then measured for fluorescence intensity (E_x_/E_m_ = 405 nm/540 nm) on a ViewLux plate reader to determine cell viability. Directly after this fluorescence reading, 4 µL of ONE-Glo™ Luciferase reagent (Promega) was added to each well using the BioRAPTR FRD™, followed by a 30-minute incubation (RT) and read for luminescence intensity on a ViewLux detector.

Activity in the agonist assay was initially defined as a sample yielding a CRC of 1.1, 1.2, 2.1, or 2.2. Activity in the antagonist assay was initially defined as a sample yielding a CRC of -1.1, -1.2, -2.1, -2.2 in the luciferase assay and selected (not exhibiting cytotoxic effects) if it has an IC_50_ ratio of >5 vs. the CellTiter-fluor (CTF) cell viability assay (if activity was observed). CTF assay was not used for agonist assays.

### P-gp assays

Assay conditions were adopted from previously described methods.^40^ The HeLa-derivative cell line KB-3-1 and its colchicine-selected, P-gp-overexpressing subline KB-8-5-11 were maintained in DMEM (Thermofisher catalog #11965) with 10% FBS and 1% penicillin/streptomycin at 37°C in 5% CO_2_. For KB-8-5-11 cells, colchicine was added to the medium at a concentration of 100 ng/ml (after initial cell attachment). For the screen, assay medium was identical to culture medium except for KB-8-5-11, where colchicine was excluded from the medium. Briefly, 5 µL of cells at 1 × 10^5^ cells/mL (500 cells/well) were dispensed into a 1536-well white solid-bottom tissue treated plate (Greiner Bio-One) using a Multidrop Combi dispenser. Plates were incubated at 37°C, 95% humidity, and 5% CO_2_ ∼4 hours to allow for cell attachment. Compounds and intraplate control (Bortezomib; final concentration range 228 pM – 7.5 µM) were transferred (16 nL) via pintool (Wako Automation), for a final concentration of 15.6 nM – 30.0 µM (most substances) and incubated for ∼72 hours at 37°C, 95% humidity, and 5% CO_2_. Plates were then dispensed with 2.5 µL of CellTiter-Glo®, centrifuged for 15 seconds at 1,000 RPM’s, followed by a 30-minute incubation (RT) and read for luminescence intensity on a ViewLux detector.

### HepG2 assay

Assay conditions for multiplexed viability output were adopted from previously described methods.^111^ HepG2 cell line purchased from ATCC were maintained in DMEM (Thermofisher catalog #11965) with 10% FBS and 1% penicillin/streptomycin at 37°C in 5% CO_2_. Cultures were confirmed to be free of mycoplasma infection using the MycoAlert Mycoplasma Detection Kit (Lonza, Walkersville, MD). Briefly, 5 µL of HepG2 cells at 2 × 10^5^ cells/mL (1,000 cells/well) were dispensed into a 1536-well white solid-bottom tissue treated plate (Greiner Bio-One) using a Multidrop Combi dispenser (Thermo Scientific). Plates were incubated at 37°C, 95% humidity, and 5% CO_2_ for ∼4 hours to allow for cell attachment. Compounds were transferred (16 nL) via pintool (Wako Automation), for a final concentration range for most substances of 15.6 nM – 31.3 µM. Plates were incubated for ∼24 hours, followed by a 1 µL addition of GF-AFC substrate (final concentration 25 µM; MP Biomedicals) and incubated at 37°C, 95% humidity, and 5% CO_2_ for 30 minutes. After incubation, plates were read for fluorescence intensity (E_x_/E_m_ = 380 nm/510 nm) on an EnVision detector (PerkinElmer, Shelton, Connecticut). Once the detection for fluorescence was completed, plates were dispensed with 3 µL of CellTiter-Glo® (Promega, Madison, WI), centrifuged for 15 seconds at 1,000 RPM’s, followed by a RT incubation for 30 minutes. Plates were then read for luminescence intensity on a ViewLux detector.

### qHTS data analysis

Data from each assay were normalized plate-wise to corresponding intra-plate controls (neutral control DMSO and positive control as noted) as described previously.^112^ The same controls were also used for the calculation of the Z’ factor, a measure of assay quality control, as previously described.^20^ Percent activity and concentration-response curves were fitted, classified, and IC_50_ determined using in-house software (http://tripod.nih.gov/curvefit/) and as previously described.^113,114^ Otherwise, concentration–response curves were fitted and IC_50_ values were calculated using Prism software (version 8, GraphPad Software, Inc. San Diego, CA), sigmoidal dose-response (variable slope). Minimum Significant Ratio (MSR), a statistical parameter that characterizes the reproducibility of potency estimates from *in vitro* concentration-response (CRC) assays,^24^ was used to assess the performance of our intraplate controls. The chemical structures were standardized using the LyChI (Layered Chemical Identifier) program (version 20141028, https://github.com/ncats/lychi).^115^ We used the Palantir Technologies (San Francisco, CA) data integration platform, which is configured to ingest all HTS results generated at NCATS and harmonized this data with other sources such as ChEMBL and OrthoMCL. All qHTS screening results are publicly available at PubChem (https://pubchem.ncbi.nlm.nih.gov/source/NCGC; AID).

### Cluster analysis

Quantitative high-throughput screening (qHTS) results obtained from nine assays for compounds from DSNP and TCM libraries, were converted into binary values using following criteria: i) compound considered to be active if curve class is negative and efficacy is more than 50%; ii) all compounds which are not passed that criteria were considered as inactive. The obtained binary activity values (active = 1 and inactive = 0) from nine assays which characterize each compound in DSNP and TCM sets (so called activity profile) were used as descriptors for clustering analysis. The clustering analysis were performed using hierarchical clustering method implemented in the KNIME analytic platform (https://www.knime.org/; accessed Feb 18, 2016). Cluster assignment was conducted using normalized Euclidian distance with threshold of 0.5, which grouped compounds together with approximately 50% similarity between them. After that compound’s activity values (active = 1 and inactive = 0) were averaging within the corresponding clusters for each particular assay and used for construction of cluster heat map utilizing Seaborn, a Python data visualization library (seaborn: statistical data visualization — seaborn 0.11.0 documentation; https://seaborn.pydata.org/; accessed Nov 4, 2020).

### Principal Component Analysis (PCA)

To analyze the compounds distribution of DSNP and TCM libraries, their biological activity profiles were used as descriptors to perform principal component analysis (PCA). Biological activity profile was calculated in the same way as described in the cluster analysis section and were used since chemical structures of some mixtures were unknown. PCA analysis was conducted using the KNIME analytic platform, resulted in utilization of 2 principal components with 76% information preservation. Thus, each compound was mapped into two-dimensional space based on two principal components values. To represent reactivity/promiscuity of compounds across all assays, we summarized binary values from all assays for each compound and used it as dot size in 2 dimensional PCA map.

### Activity Frequency Analysis

Average activity frequency of a compound was calculated as percent of assays, in which the compound is active. Distribution of activity frequency of compounds within the clusters defined in the Cluster analysis section were calculated and visualized using whisker plot of the seaborn python data visualization library to reveal the most and less reactive/promiscuous clusters.

### Activity Profile Analysis

Analysis of compound concentration–response data was performed as previously described.^116^ Briefly, concentration–response titration points for each compound were fitted to a four-parameter Hill equation. Compounds were designated as Class 1–4 according to the type of concentration–response curve observed.^113,116^ Each curve class was then converted to a curve rank^116^ such that more potent and efficacious compounds with higher quality curves were assigned a higher rank. The curve rank is a value ranging from -9 to +9, with -9 to -1 indicating inhibitory ability and 1 to 9 indicating activating ability, and 0 meaning inactive.^116,117^ Curve ranks should be viewed as qualitative descriptors of the concentration response profile of the compound. Hierarchical clustering was performed on the NPC together with the DSHEA/TCM libraries based on similarity in their compound activity profiles (measured by curve rank) across the 15 assays using the UPGMA method in TIBCO® Spotfire® version 7.11.1, resulting in 259 clusters and 141 singletons. Compound mode of action (MOA) annotations were retrieved from the Medical Subject Headings (MeSH) (http://www.ncbi.nlm.nih.gov/mesh) database (pharmacological action (PA) terms) and the KEGG Drug database (https://www.genome.jp/kegg/drug/). Each cluster was evaluated for enrichment of MOA terms using the Fisher’s exact test. Heat maps and PCA plots were generated using TIBCO® Spotfire® version 7.11.1.

## Figure Legends

**Figure S1.**
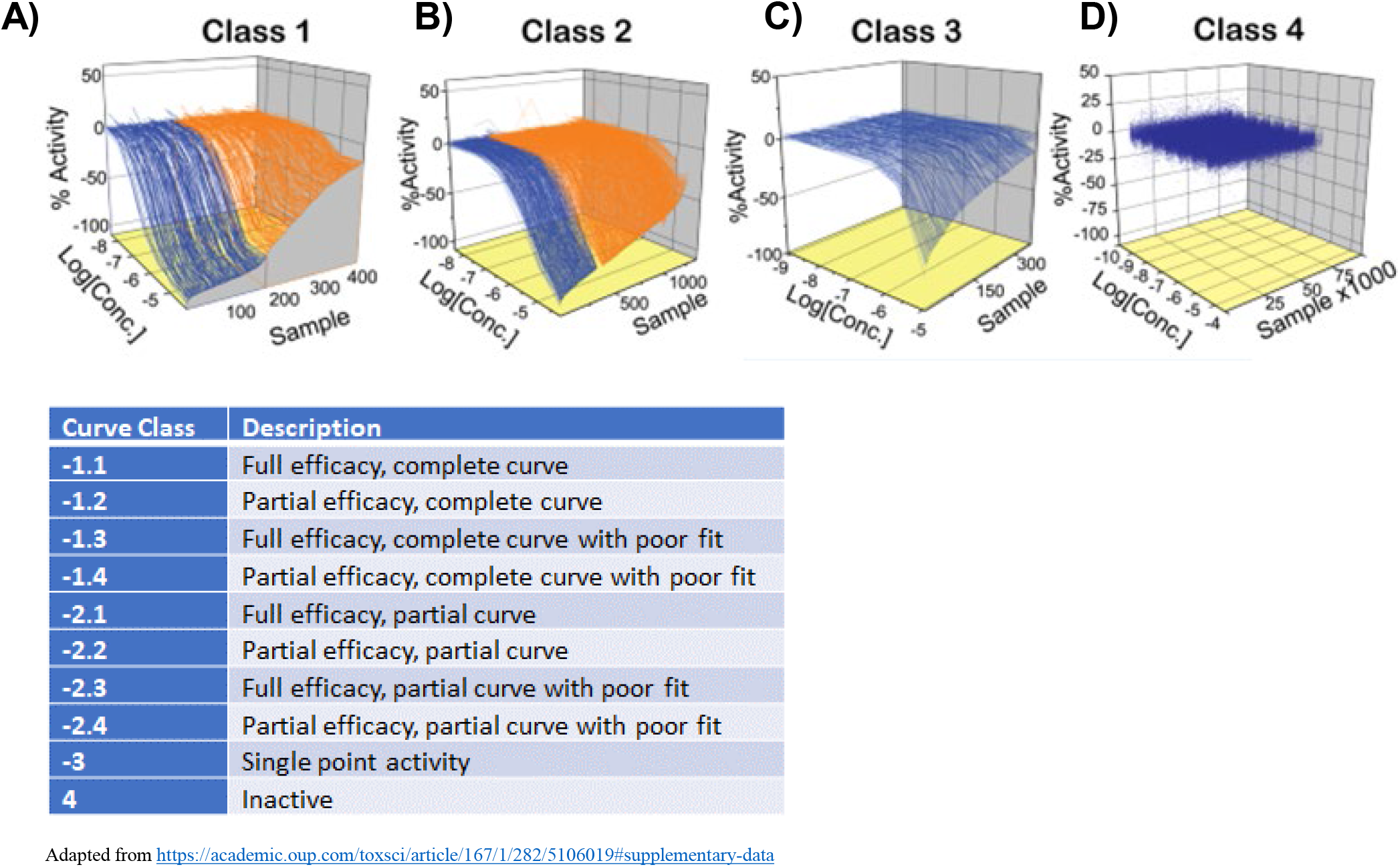
Example curves highlighting qHTS curve classification criteria. Lines connecting titration data corresponding to inhibitory compounds are shown. (a) Classes 1.1 (blue; >80% efficacy) and 1.2 (orange; ≤80% efficacy) inhibitors display full and partial activity, respectively, with r2 ≥0.9. (b) Incomplete curves for inhibitors having AC50 values within and beyond the tested titration range are Classes 2.1 (blue; >80% efficacy, r2 >0.9) and 2.2 (orange; ≤80% efficacy, r2 <0.9), respectively. (c) Incomplete inhibitory (blue) curves that show weak activity and poor fits are Class 3. (d) inactive compounds are Class 4.

**Figure S2.**
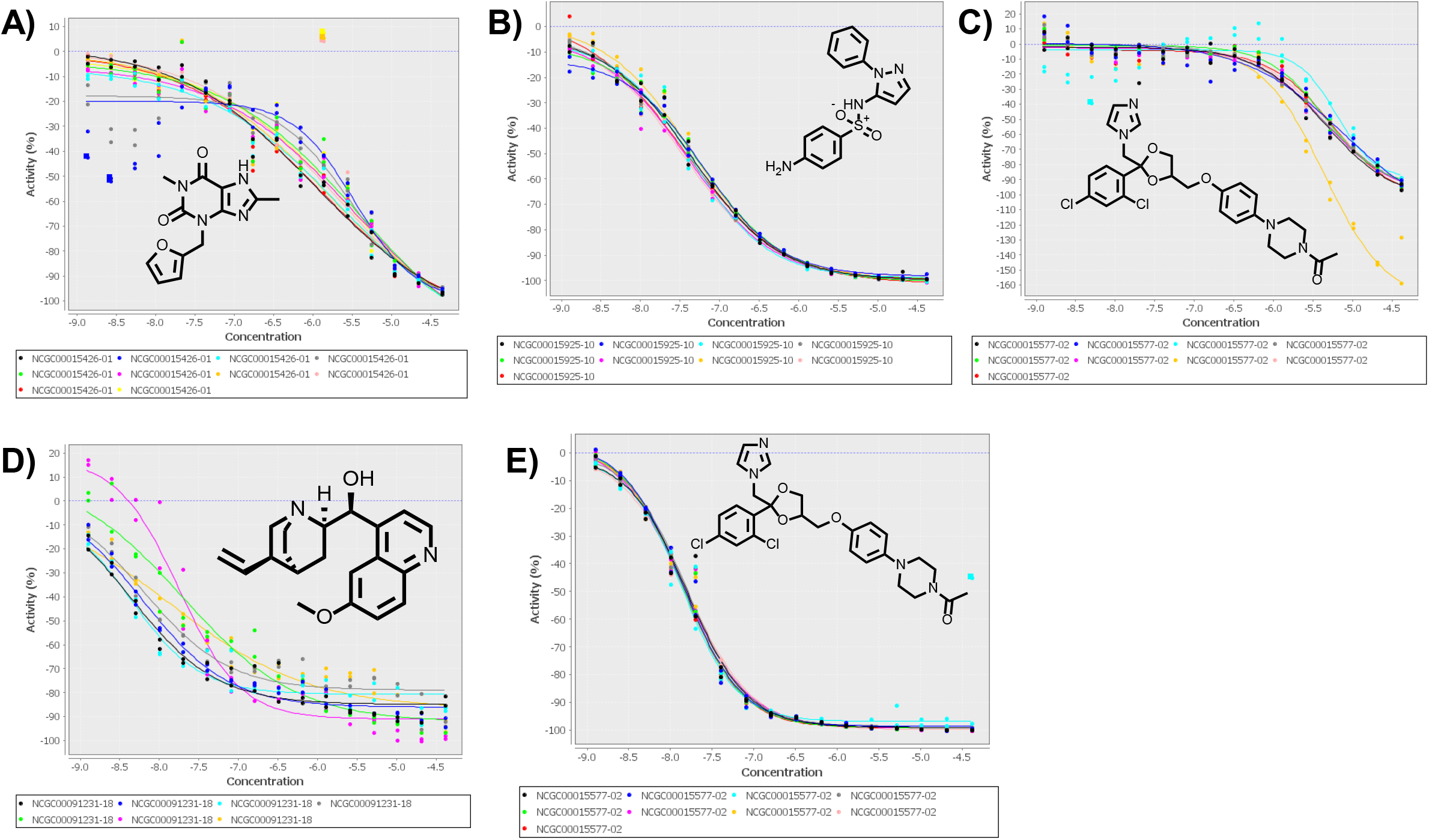
The intraplate controls Furafylline (CYP1A2), Sulfaphenazole (CYP2C9), Ketoconazole (CYP2C19), Quinidine (CYP2D6), and Ketoconazole (CYP3A4) also performed well, with mean IC_50_’s [µM] of 2.60 (MSR = 3.57), 0.0448 (MSR = 1.78), 5.17 (MSR = 1.82), 0.103 (MSR = 5.98), and 0.0145 (MSR = 1.18), respectively

**Figure S3.**
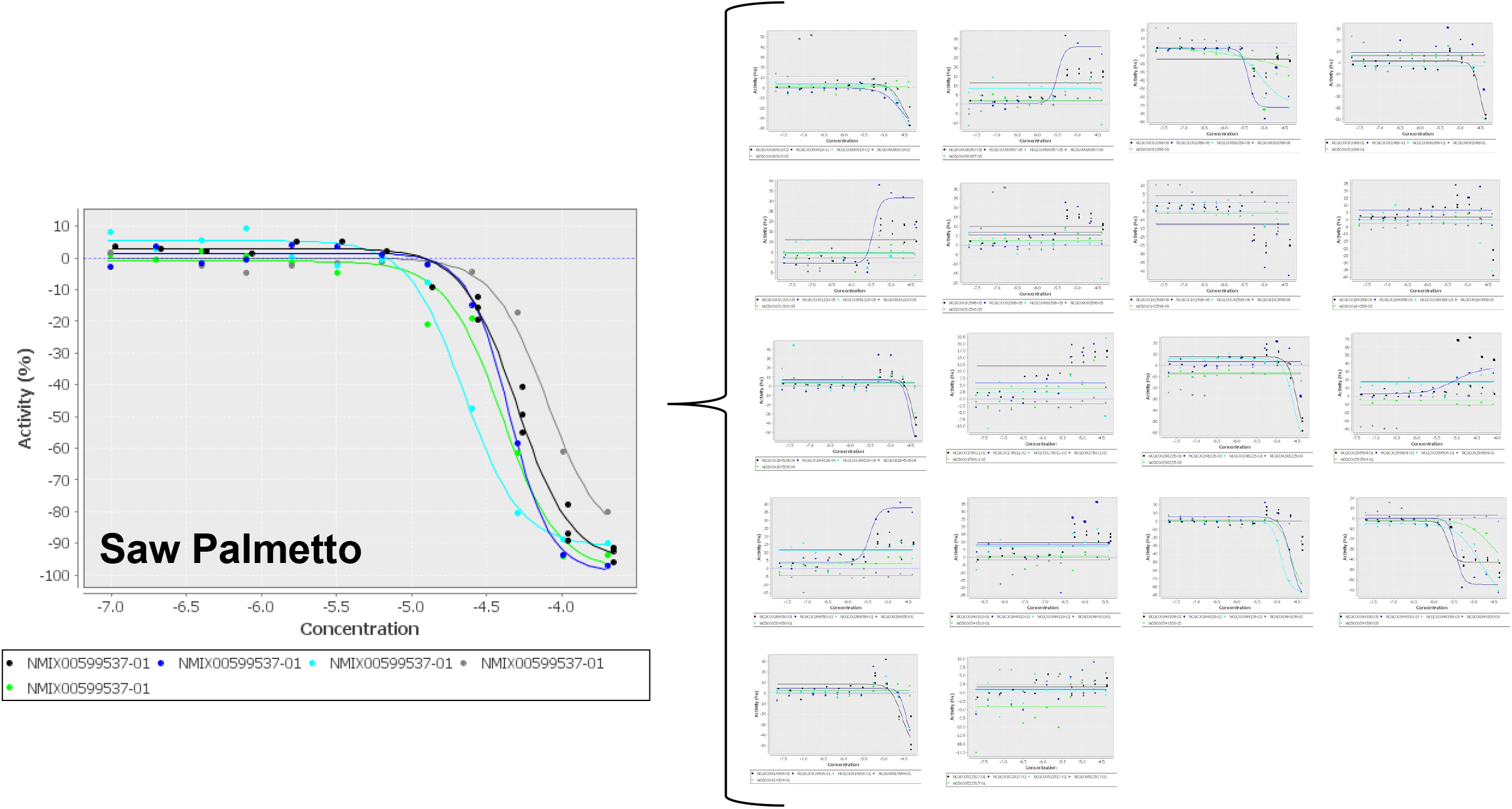
Saw Palmetto (NMIX00599537) has associated 18 constituents (based on GSRS database). Saw Palmetto was pan-active with an IC50 range of 22.4 – 79.4 µg/mL, three compounds, Linoleic acid (NCGC00344326), Oleic acid (NCGC00344330), and Lauric acid (NCGC00090919), appear to be responsible for the majority of its P450 activity.

**Figure S4.**
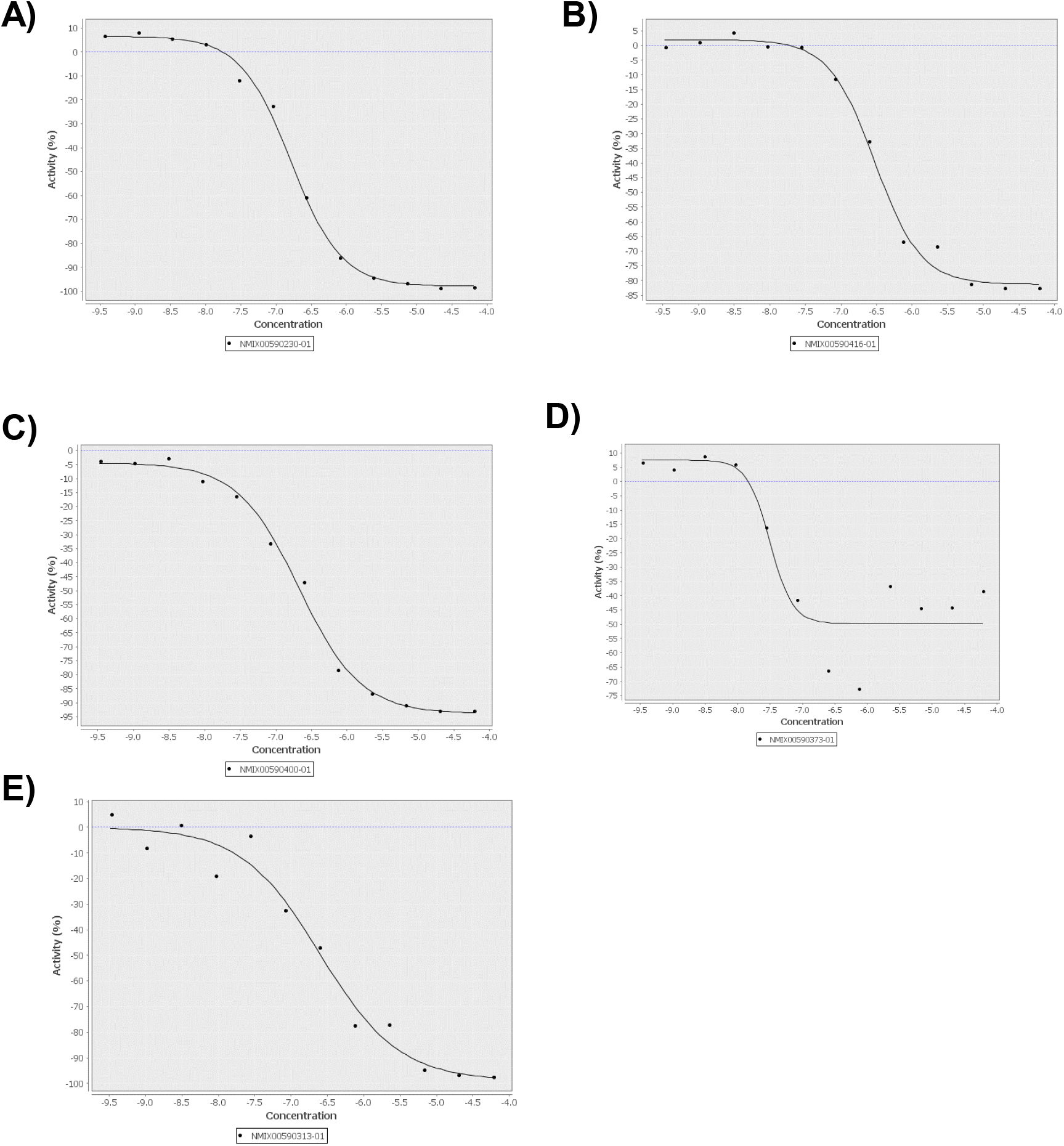
The most potent samples were all organic extracts (IC_50_ values < 0.3 μg/mL) for CYP’s 1A2, 2C9, 2C19, 2D6, and 3A4 were derived from Angelica dahurica (root of the Holy Ghost, NMIX00590230), Sophora flavescens (Ku Shen, NMIX00590416), Juncus effusus (Soft Rush; NMIX00590400), Corydalis yanhusuo (NMIX00590373) and Piper nigrum (Black pepper, NMIX00590313), respectively

**Figure S5.**
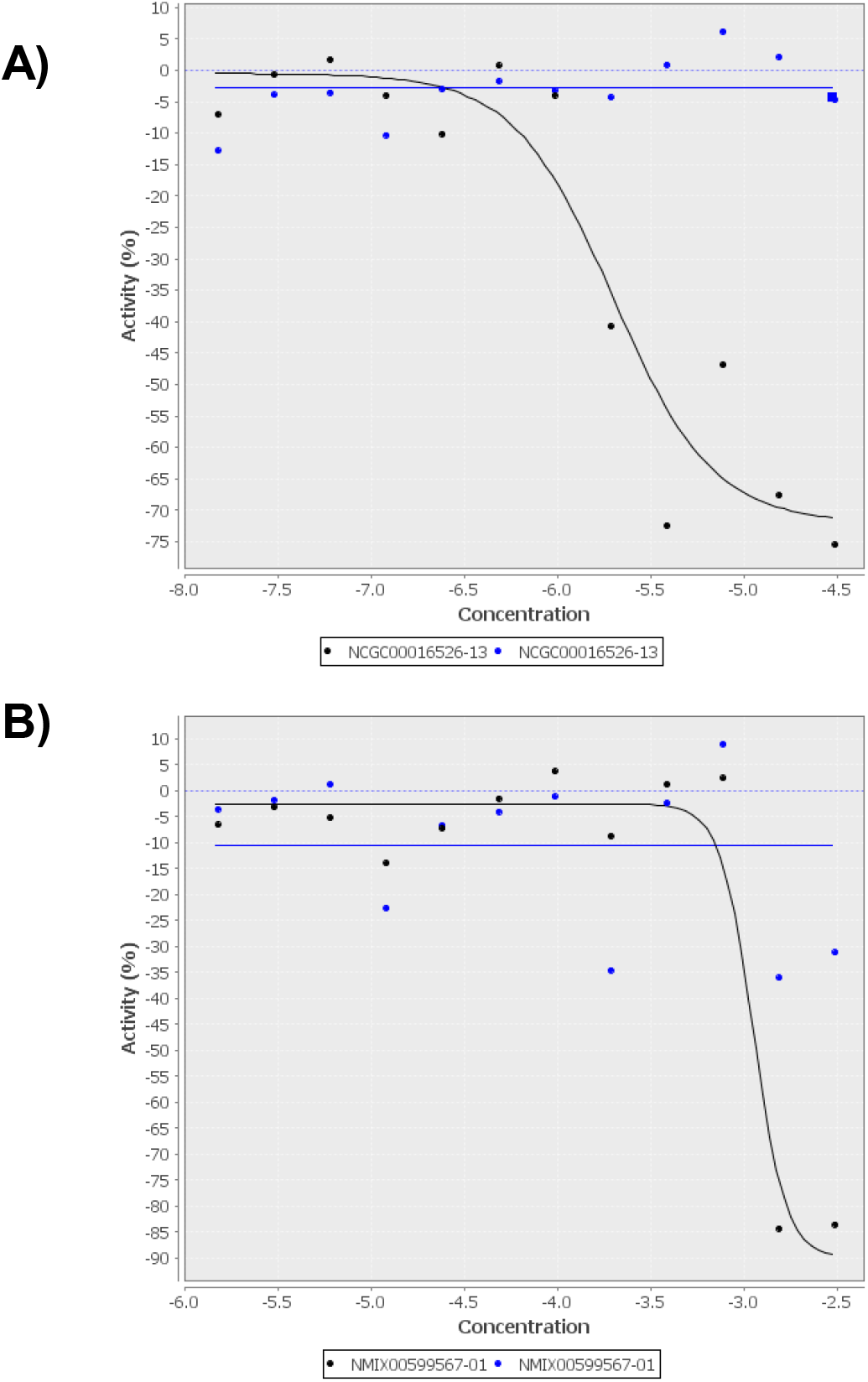
The supplement Berberine (Panel A) and botanical Flaxseed Oil (Panel B) exhibited cytotoxicity in the KB-3-1 cell line with IC50’s of 2.0 µM and 1,122 μg/mL, respectively) and were inactive in the KB-8-5-11 cell line (blue).

**Figure S6.**
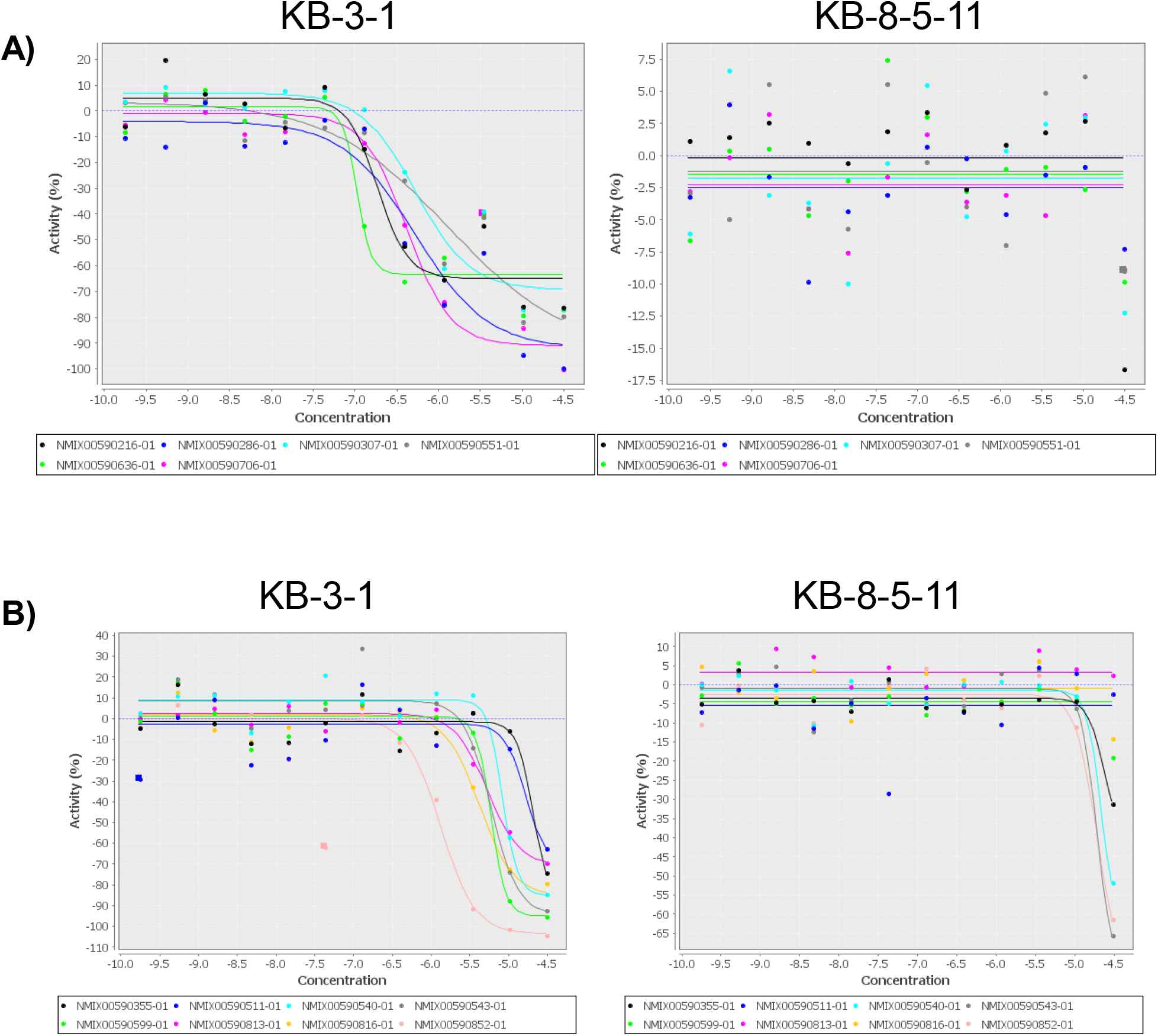
Coptis chinensis (Chinese Goldthread) cellular activity shown in Panel A; Brucea javanica (Macassar kernels) cell differential activity in Panel B.

